# Partitioning gene-level contributions to complex-trait heritability by allele frequency identifies disease-relevant genes

**DOI:** 10.1101/2021.08.17.456722

**Authors:** Kathryn S. Burch, Kangcheng Hou, Yi Ding, Yifei Wang, Steven Gazal, Huwenbo Shi, Bogdan Pasaniuc

## Abstract

Recent works have shown that SNP-heritability—which is dominated by low-effect common variants—may not be the most relevant quantity for localizing high-effect/critical disease genes. Here, we introduce methods to estimate the proportion of phenotypic variance explained by a given assignment of SNPs to a single gene (*genelevel heritability*). We partition gene-level heritability across minor allele frequency (MAF) classes to find genes whose gene-level heritability is explained exclusively by “low-frequency/rare” variants (0.5% ≤ MAF < 1%). Applying our method to ~17K protein-coding genes and 25 quantitative traits in the UK Biobank (N=290K), we find that, on average across traits, ~2.5% of nonzero-heritability genes have a rare-variant component, and only ~0.8% (370 gene-trait pairs) have heritability exclusively from rare variants. Of these 370 gene-trait pairs, 37% were not detected by existing gene-level association testing methods, likely because existing methods combine signal from all variants in a region irrespective of MAF class. Many of the additional genes we identify are implicated in phenotypically related Mendelian disorders or congenital developmental disorders, providing further evidence of their trait-relevance. Notably, the rare-variant component of gene-level heritability exhibits trends different from those of common-variant gene-level heritability. For example, while total gene-level heritability increases with gene length, the rare-variant component is significantly larger among shorter genes; the cumulative distributions of gene-level heritability also vary across traits and reveal differences in the relative contributions of rare/common variants to overall gene-level polygenicity. We conclude that the proportion of gene-level heritability attributable to low-frequency/rare variation can yield novel insights into complex-trait genetic architecture.

## Introduction

Since the vast majority of risk variants identified through genome-wide association studies (GWAS) are located in noncoding regions, the genes and pathways driving complex traits are largely unknown^1–3^. For most complex traits, fundamental characteristics of genetic architecture—for example, the number of variants/genes with nonzero effects (polygenicity), the number of genes regulated by local versus distal variants, and the relative contributions of rare versus common variants to gene expression and phenotype—remain actively debated^4–14^.

That complex-trait SNP-heritability is enriched in regulatory regions is well established^1,15–17^. However, since SNP-heritability is overwhelmingly driven by common variants of low effect—individual rare variants with large per-allele effects contribute very little to population-level phenotypic variance^18,19^—whether the largest heritability enrichments localize the most clinically relevant regions and/or genes for a trait is unclear. For example, a recent study estimates that the majority of complex-trait SNP-heritability mediated via the *cis*-genetic component of expression is explained by genes that individually have low *cis*-heritability of expression^20^. In addition, despite the inherent complexity of the biological processes driving complex traits, there is growing evidence that extreme complex-trait polygenicity may be explained in large part by negative/stabilizing selection, which purges high-effect alleles from the population, producing the remarkably even distribution of SNP-heritability among common variants genome-wide (the so-called “flattening” hypothesis)^21,22^. If the most critical genes for a trait are not necessarily localized by enrichments of total heritability^20,21,23,24^, the open question of how to identify target genes using heritability enrichments or overlaps between GWAS and expression quantitative trait loci^25,26^ becomes even murkier. Gene-based association tests that aggregate signal from multiple rare variants—for example, burden tests and sequence-based association tests (SKAT)—can increase power under different genetic-architecture scenarios^27–36^. However, such methods are generally designed to test for only rare-variant association or the combined effects of common and rare variants, and thus are not ideal for parsing the relative contributions of rare/common variants to the heritability of a single gene.

Here, we propose an approach to estimate the relative heritability contributions of common, low-frequency, and rare variants to a quantity we call *gene-level heritability* 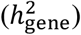, defined as the proportion of phenotypic variance explained by the additive effects of a given set of variants assigned to a gene of interest. While the method itself is general and can be applied to any small annotation of interest (see Discussion), our goal in this work is to use MAF-partitioned gene-level heritability estimates to identify disease-relevant genes, which may have different relative contributions to heritability across MAF classes. The key challenge in estimating gene-level heritability lies in the *uncertainty* about which variants are causal and what their causal effect sizes are; such uncertainty in fine-mapping increases as the strength of LD in the region increases and as GWAS sample size decreases^37^. Consider a toy example in which a variant in the gene of interest is in perfect LD (LD=1) with a second variant adjacent to the gene, the observed data are GWAS marginal association statistics and LD for the region (Figure 1a). Without additional information, it is impossible to definitively elucidate the underlying causal configuration.

**Figure 1.**
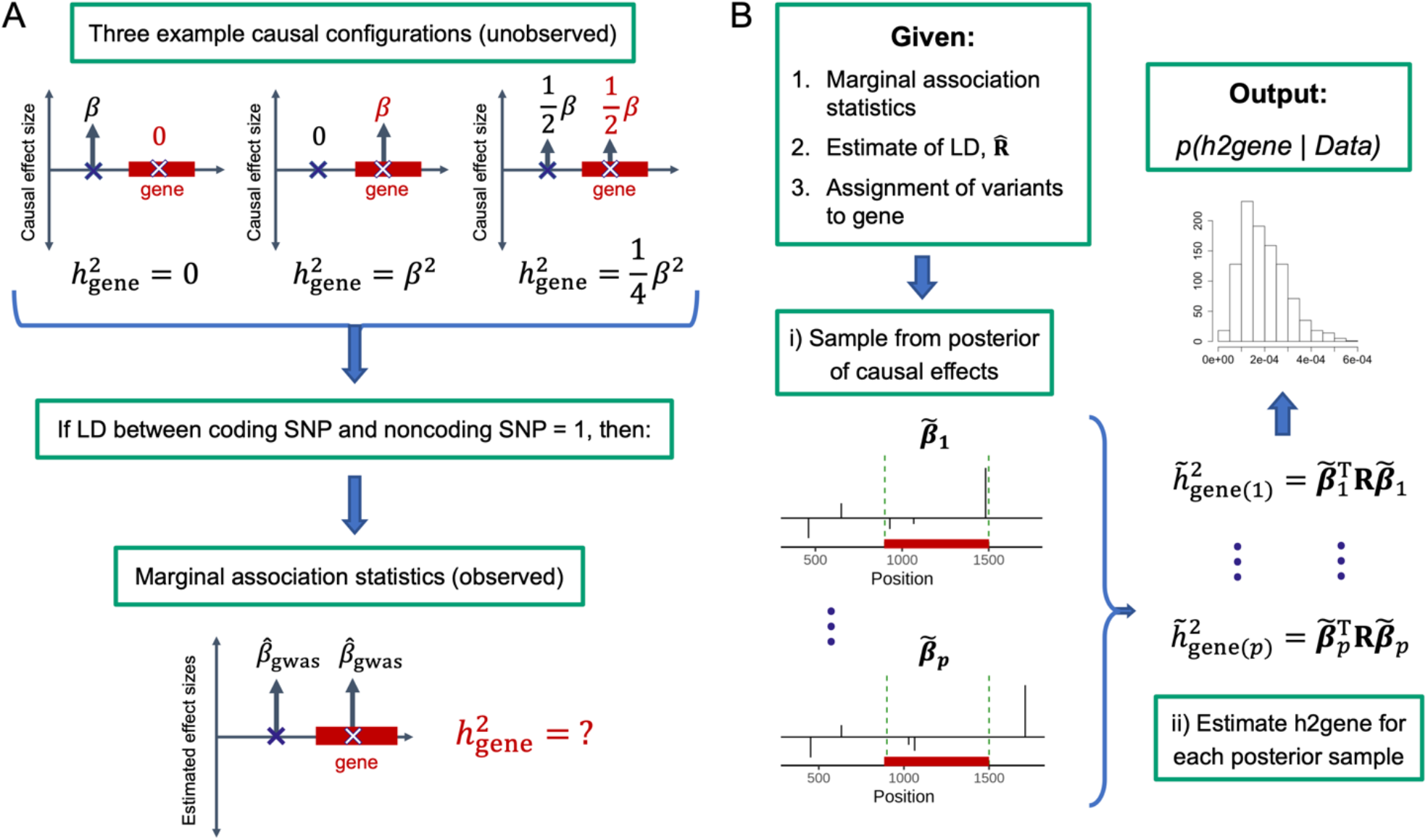
Overview. (A) Toy example with two variants, one of which is assigned to the gene of interest. The top row depicts 3 example causal configurations corresponding to 3 different gene-level heritabilities (0, *β*^2^, and *β*^2^/4). Since the variants in are in perfect LD, all 3 causal configurations yield the same expected marginal association statistics. (B) Given marginal association statistics, an estimate of LD, and an assignment of variants to the gene of interest, our approach involves i) sampling from the posterior of the causal effect sizes (assuming a sparse prior) to capture our uncertainty about which variants are causal, and then ii) estimating gene-level heritability for each posterior sample to approximate the posterior distribution of gene-level heritability.

Even if the LD between the variants is 0.9 instead of 1, if this GWAS has 90% power to identify the associated region, to correctly reject the null hypothesis for the non-causal variant would require a sample size ≥ 4x larger than that of the original GWAS^37^. Since each causal configuration can yield a different gene-level heritability (with or without MAF-partitioning), randomly selecting one possible configuration (e.g., using variable selection methods such as the Lasso^38^) can yield inaccurate/misleading estimates. As an alternative approach, methods for partitioning genome-wide SNP-heritability across MAF bins can be employed. However, such methods are also ill-suited to our goals as they make distributional assumptions on the causal effects which (i) limit power to detect enrichment in small categories of variants (< 1% of the genome) and/or (ii) may not apply equally to rare and common variants^15,17,39–43^. Estimators for the SNP-heritability of a single region (“regional SNP-heritability”) yield inflated estimates if any variants in the region of interest are in LD with the adjacent regions^23,44–46^. To address the fine-mapping uncertainty, we seek to propagate the uncertainty about which variants are causal to infer the posterior distribution over the entire gene of interest. Given GWAS summary statistics and estimates of LD, we sample from the posterior distribution of the causal effect sizes within a probabilistic fine-mapping framework^47^ and use the posterior samples to approximate the posterior distribution of gene-level heritability, thus capturing uncertainty in the causal effects (Figure 1b). From the full posterior distribution of gene-level heritability, one can compute various summary statistics of interest for each gene. We report the posterior mean, which we denote 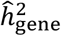, and *ρ*-level credible intervals, or *ρ*-CI, defined as the central interval containing the true gene-level heritability with probability *ρ* (Material and Methods).

We confirm in simulations that accounting for uncertainty in the estimated causal effects significantly reduces the bias of 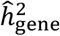. Although the corresponding *ρ*-CIs are not perfectly calibrated—for example, at *ρ* = 0.9, about 70% of credible intervals overlap 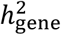—among the true causal genes, any mis-calibrated CIs overwhelmingly tend to underestimate rather than overestimate 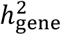. Both 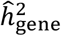 and *ρ*-CIs are robust to parameters such as causal effect sizes, gene length, allele frequencies of causal variants, and the strength of local LD. Assuming that total genelevel heritability can be expressed as 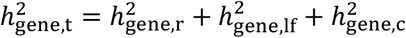, where each term refers to the component of 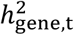 explained by rare (0.5% ≤ MAF < 1%), low-frequency (1% ≤ MAF < 5%), and common (MAF ≥ 5%) variants, respectively, we apply the same approach to estimate the posterior distributions of 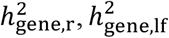, and 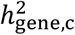 and observe similar trends and levels of accuracy (we note that there are many definitions of “rare” in the literature, and that we use 0.5% ≤ MAF < 1% because we analyze imputed genotypes).

Applying our approach to estimate gene-level heritability for 17,436 genes and 25 quantitative traits in the UK Biobank^48^ (N=290K self-reported “white British”, MAF > 0.5%), we find that 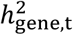 is indeed dominated by 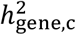. Among genes with 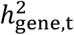 90%-CI > 0 (“nonzero-heritability genes”) for a given trait, 92% (s.d. 1%) have nonzero common-variant heritability, and 76% (s.d. 1%) have nonzero heritability exclusively from common variants (i.e. 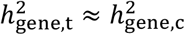). In contrast, only 2.5% (s.d. 0.6%) of nonzero-heritability genes, averaged across traits, have nonzero rare-variant heritability, and 0.8% (s.d. 0.4%) have nonzero heritability exclusively from rare variants 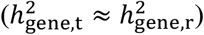. As a sanity check, we confirm that Mendelian-disorder genes from OMIM^49^, genes intolerant to loss of function (LoF) variants^50^, and a set of FDA-approved drug targets for 30 immune-related traits^51^ have elevated estimates of all four heritability quantities (total, common, low-frequency, and rare). Among the 0.8% with 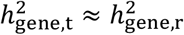 (370 gene-trait pairs in total), we identify many examples of disease genes with known roles in phenotypically similar Mendelian disorders and other congenital growth and developmental disorders. 37% of the 370 gene-trait pairs were not identified by existing methods for gene-level association testing, likely because existing methods have low power to detect genes containing only rare variants of moderate or low effect. We observe an overrepresentation of LoF-intolerant genes, but not Mendelian-disorder genes, among the 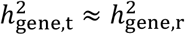 genes. Using gene-level heritability estimates to further explore genetic architecture reveals notable differences between total/rare-variant gene-level heritability; for example, while total/common-variant gene-level heritability increases with gene length, we observe a clear inverse relationship between the rare-variant component and gene length.

Taken together, our results show that the low-frequency/rare-variant component of total gene-level heritability is useful for identifying narrow sets of high-impact genes that are not necessarily located in regions enriched with common-variant heritability. Our results are also consistent with the hypothesis that a sizable amount of complex-trait variation is driven by dysregulation of genes that—if completely disrupted—cause phenotypically similar monogenic disorders and/or systemic congenital and developmental disorders^52^. Since some high-impact genes are disrupted/dysregulated by a combination of common and rare variants, we conclude that 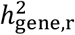 should be considered alongside common-variant heritability enrichments if one is interested in identifying high-impact disease genes under different degrees of selection. While we restrict our analyses to genes (± 10-kb window), our method is general and can thus be applied to any small annotation of interest (e.g., enhancers, a set of genes involved in a pathway, a set of putative causal variants).

## Results

### Overview of the Methods

We propose a general approach for estimating the heritability explained by a given set of variants and assess its utility in estimating gene-level heritability. Given an assignment of *m* variants to a gene *g* of interest, total genelevel heritability is defined as 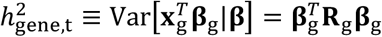, where **β**_g_ is the *m* × 1 vector of unknown causal effect sizes and **R**_g_ is the *m* × *m* LD for SNPs in the gene (Material and Methods). Our goal in this work is to estimate a *distribution* over 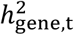 that captures uncertainty in the causal effects that arises from LD (Figure 1a). To this end, we adopt a probabilistic fine-mapping framework^46,47^ which assumes a sparse prior on the causal effect sizes in the LD block containing gene *g* and infers the posterior distribution of the causal effect sizes, 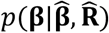, where 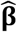 is the vector of estimated marginal effects from GWAS and 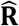 is an estimate of LD. We sample from the posterior of **β** to approximate to the posterior of 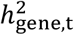 (Figure 1b, Material and Methods). For each gene, we report the estimated posterior mean, denoted 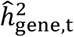, and *ρ*-level credible intervals (*ρ*-CI), defined as the central interval that contains the true gene-level heritability with probability *ρ* ∈ [0, 1]. Whereas previous works applied similar approaches to generate credible sets of causal variants^47^ or to estimate regional SNP-heritability of LD blocks^46^, our goal in this work is to estimate the heritability explained by any arbitrary (not necessarily contiguous) set of variants much smaller than an LD block. This allows us to partition by minor allele frequency (MAF) bins under the assumption that 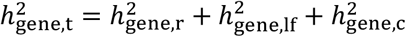, where the subscripts represent the rare (0.5% ≤ MAF < 1%), low-frequency (1% ≤ MAF < 5%), and common (MAF ≥ 5%) variants assigned to the gene. (We note that, while there are many definitions of “rare” in the literature, we threshold at MAF ≥ 0.5% because we want to reduce potential noise from imputation; see Discussion for details.)

### Accuracy of gene-level heritability estimates in simulations

We perform simulations starting from real imputed genotypes of N=290,273 “unrelated white British” individuals in the UK Biobank (chromosome 1, MAF > 0.5%, M=200,235 variants, 1,083 genes; Material and Methods). In all simulations, the estimand of interest (gene-level heritability, 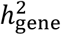) is the proportion of phenotypic variance explained by the variants in the gene body, as well as the MAF-partitioned counterpart. We note that our choice of variant assignment is arbitrary; there are many ways to assign variants to a gene, but our goal in this section is to provide a proof of concept. In brief, our simulation framework consists of three steps. First, for a given total heritability (variance explained by all *M* variants) and cumulative gene-level heritability (variance explained by all genes), we randomly select 3%, 8%, or 16% of the genes to be causal, where “causal” in this context refers to genes with 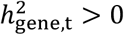. Second, for each causal gene, we draw causal variants in the gene body and within 10-kb upstream/downstream of the gene start/end positions; the purpose of the latter is to create situations where the estimated effects of variants in the region of interest are inflated in part because they tag causal effects located adjacent to the region. Third, we sample noncoding “background” causal variants from the whole chromosome with frequency *p*_causal_ = {0.001, 0.01}. Under this model, the majority of simulated gene-level heritabilities are on the order of 10^-6^ to 10^-3^ (Supplementary Figure 1), similar to what we observe in real data in subsequent sections.

Overall, the estimated posterior means of total gene-level heritability, 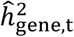, are highly concordant with the true gene-level heritabilities (Figure 2, Supplementary Figure 2). For each gene, we compute two metrics of accuracy from *s* = 30 simulation replicates: 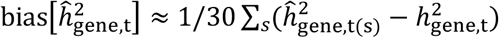, and 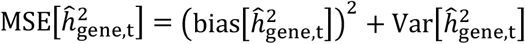 (mean squared error) (Material and Methods). As expected, MSE increases as the background polygenicity (*p*_causal_) and proportion of causal genes increase, i.e. as causal effect sizes of noncoding variants and gene-level heritabilities decrease (Supplementary Figure 3). Among the causal genes 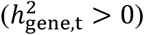, 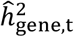 tends to underestimate 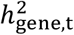, with the median bias across genes ranging from approximately 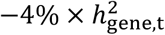 for lower polygenicities to −30% × 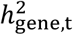 for higher polygenicities (Figure 2, Supplementary Figure 4). There is a small positive correlation between bias and gene length (average Pearson *R* = 0.05 (s.d. 0.02) across simulation setups), i.e. the estimates tend to be more downward-biased for shorter genes; average LD score and average MAF of variants in the gene have no discernible impact on accuracy (Supplementary Figures 5-8). To visualize the impact of causal-effect uncertainty on gene-level heritability estimation, we compare 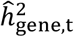 to a naive estimator that ignores LD between the gene and its adjacent regions, thus ignoring causal-effect uncertainty (Material and Methods). As expected, the naive estimator is significantly inflated; in particular, many noncausal genes have dramatically upward-biased estimates (Figure 2, Supplementary Figures 2 and 9) due to LD between variants in the gene and nearby causal variants. We benchmark the estimators for the contributions of rare, low-frequency, and common variants to total gene-level heritability and find that they perform similarly to 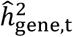 (Figure 3, Supplementary Figures 3, 4, 6-8, 10-12).

**Figure 2.**
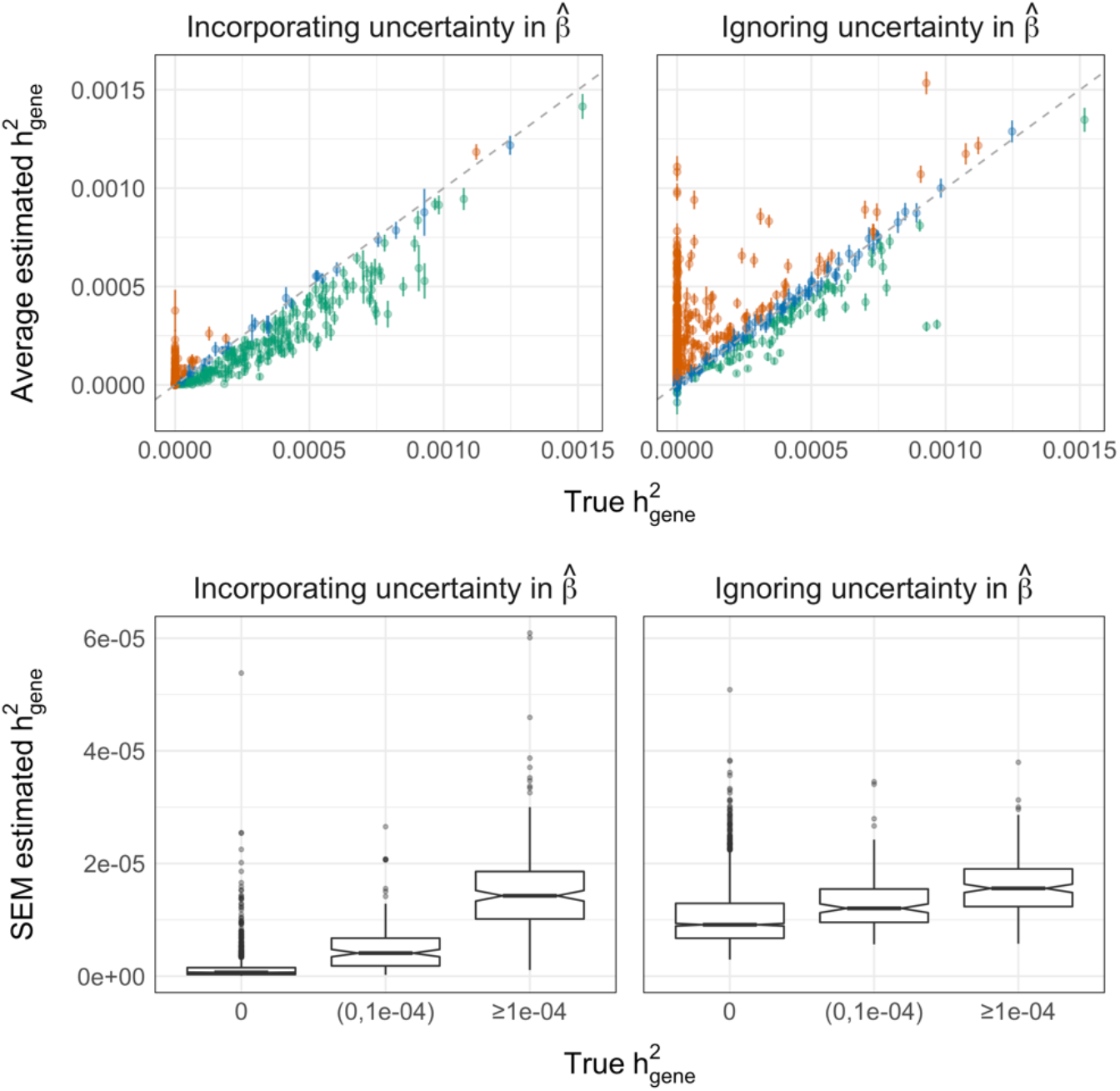
Impact of uncertainty in the estimated causal effects on gene-level heritability estimation in simulations. Chromosome 1, MAF > 0.5%, *P*_causal_=0.01, N=290K individuals, and 1,038 genes, of which 16% have nonzero gene-level heritability. Top row: each point is the average 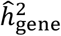 for a given gene across 30 simulation replicates; error bars mark 1.96 × standard error of the mean (SEM). Orange and green points are genes for which the estimator is significantly upward-biased and downward-biased, respectively. Bottom row: distributions of SEM with respect to gene-level heritability.

**Figure 3.**
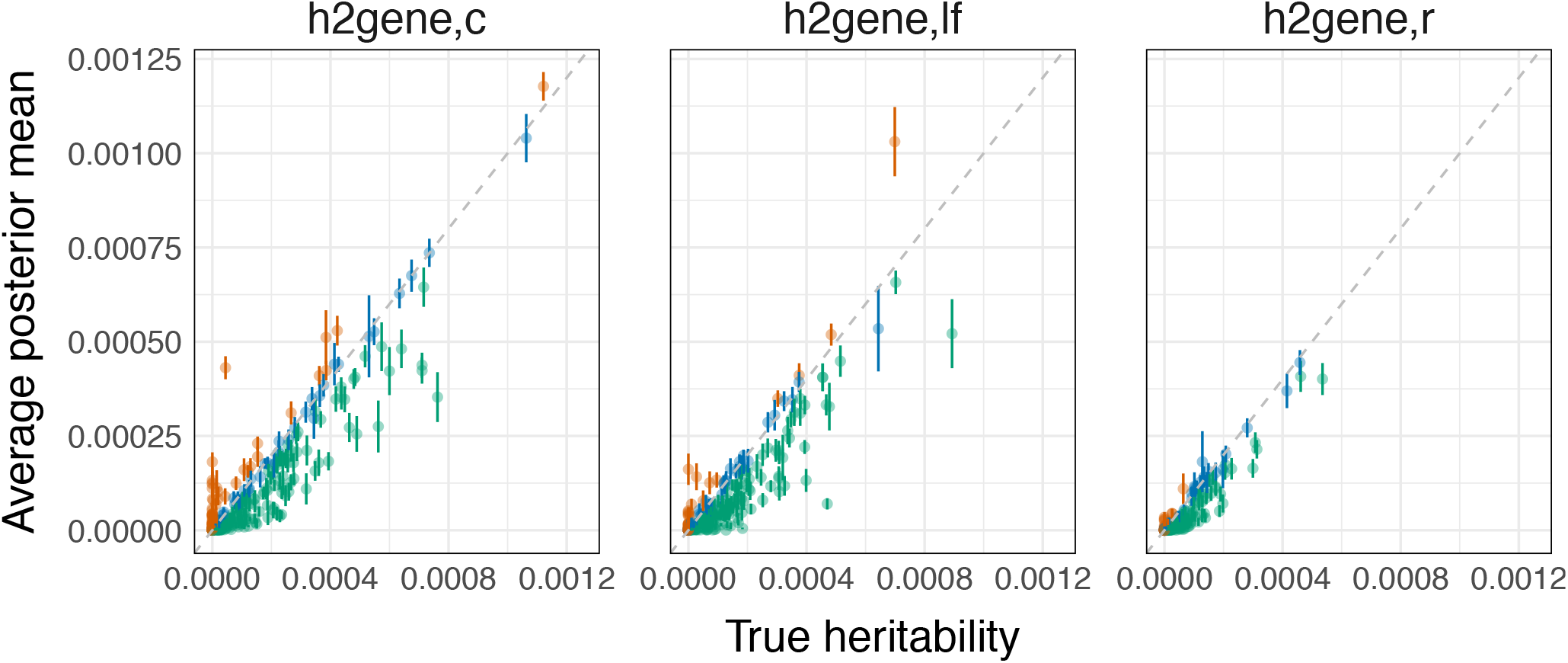
Estimates of the heritability contributions of common, low-frequency, and rare variants in simulations. Chromosome 1, MAF > 0.5%, *P*_causal_=0.01, N=290K individuals, and 1,083 genes, of which 16% have nonzero heritability. Each point is the average posterior mean for a given gene from 30 simulation replicates; error bars mark 1.96 x SEM. Orange and green points are genes for which the estimator is significantly upward-biased and downward-biased, respectively, where significance is determined by the error bars.

### Calibration of credible intervals

Calibration of *ρ*-level credible intervals (*ρ*-CIs) was assessed using “empirical coverage,” defined here as the proportion of simulation replicates in which *ρ*-CI contains the true gene-level heritability (Material and Methods). Perfect calibration of *ρ*-CI would manifest as empirical coverage equal to *ρ* for all *ρ* ∈ [0, 1]. In reality, we observe a downward bias in empirical coverage across all simulations that increases in magnitude as the proportion of causal genes increases (i.e. as per-variant causal effect sizes decrease). For example, at *ρ* = 0.9, empirical coverage ranges from approximately 0.75 when 3% of genes are causal to 0.65 when 16% are causal (Supplementary Figure 13). While downward bias in empirical coverage can be the result of *ρ*-CIs underestimating or overestimating 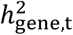, the credible intervals at *ρ* = {0.90, 0.95} tend to underestimate the true gene-level heritability (Supplementary Table 1), consistent with the downward-bias we observe in 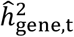 (Figure 2). For example, at *ρ* = 0.95, the proportion of true causal genes that are underestimated vs. overestimated is approximately 14% vs. 6% (when 3% of genes are causal) and 30% vs. 3.5% (when 16% of genes are causal) (Supplementary Table 1). The *ρ*-CIs for 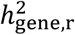 are more conservative; for the same parameters, among the genes with true 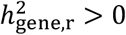, the proportions of underestimated vs overestimated genes are 38% vs. 1.5% (when 3% of genes are causal) and 45% vs. <1% (when 16% of genes are causal) (Supplementary Table 2, Supplementary Figure 14).

### Robustness to noise in estimates of LD

Finally, we assess whether 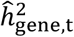 is robust to the number of individuals used to estimate LD, i.e. the sample size of the “LD panel” (Material and Methods). Compared to in-sample LD computed from the full set of individuals in the GWAS (N = 290,273), using a random subset of N={500, 1000, 2500, 5000} individuals from the original GWAS does not significantly impact the MSE of 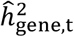 or 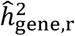 (Supplementary Figure 15). Using 90%-CIs to identify potential causal genes (i.e. 90%-CI lower bound > 0), we observe a slight increase in the false positive rate for both 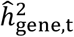 and 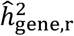 as N decreases (Supplementary Figure 16); this is accompanied by a slight increase in power for 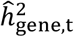 but not for 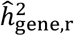 (Supplementary Figure 17). Since the N=5,000 LD panel and the full in-sample LD yield similar false positive rates for both estimators, we recommend using an in-sample LD panel of no less than 5,000 individuals (see Discussion for additional comments on LD panels).

### 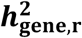 identifies genes that link complex traits to phenotypically related monogenic disorders

We estimate, and partition by MAF, the gene-level heritabilities of 17,437 genes for 25 quantitative traits in the UK Biobank (N=290,273 “unrelated white British” individuals^48^, M=5,650,812 with MAF > 0.5%, imputed data; Material and Methods). Unless otherwise stated, the quantity of interest, 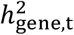, is a function of the variants located in the gene body *and* the variants located within 10-kb upstream/downstream from the gene start/end positions. A gene is classified as having “nonzero heritability” if it meets two criteria: (i) the 90%-CI for 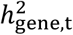 does not overlap zero and (ii) the 90%-CI for at least one MAF component (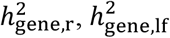, or 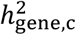) does not overlap zero. Using this definition, the number of nonzero-heritability genes ranges from 1,212 (7%) for corneal hysteresis to 2,469 (14%) for height (Table 1). Most of the estimated posterior means for these genes lie between 10^-6^ and 10^-4^ (Figure 4).

**Figure 4.**
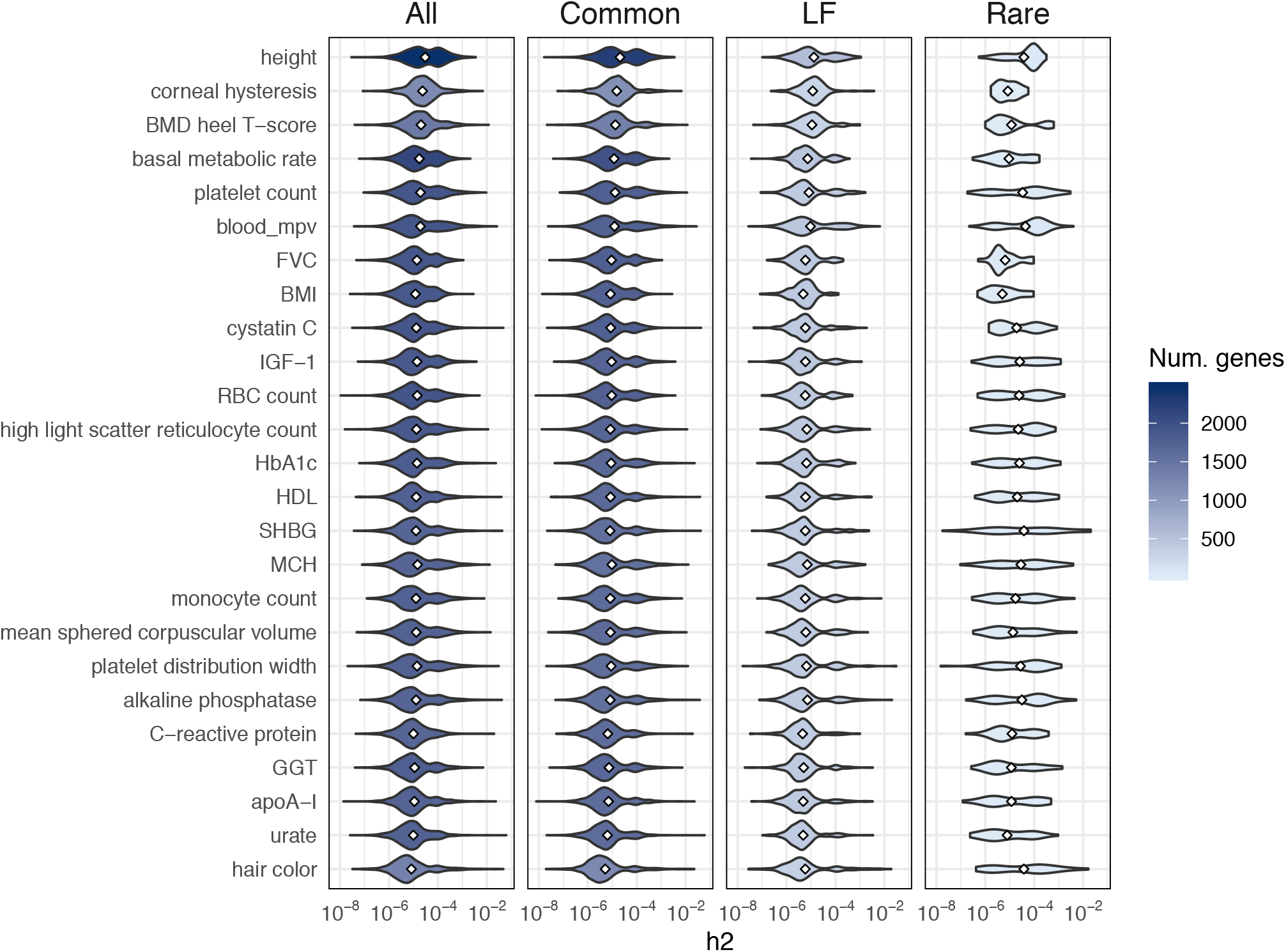
Distributions of h2 estimates for 25 traits. Each violin plot is the distribution of posterior mean estimates for genes with 90%-CI > 0 for one trait. The shading scales with the number of genes in the violin plot.

**Table 1.**
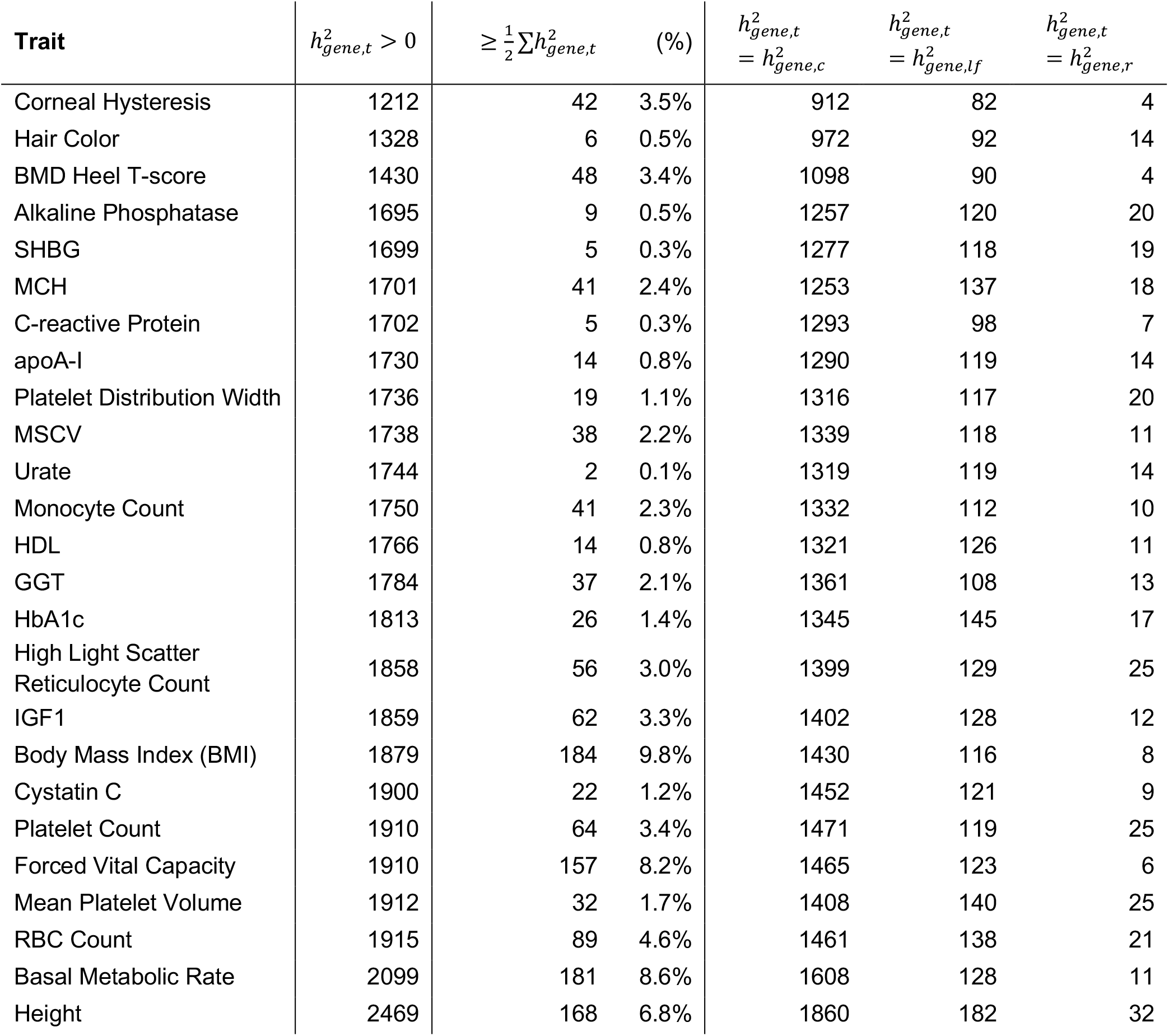
Summary of nonzero-heritability genes (90%-CI) for 25 quantitative traits. Columns 1-4: complex trait; total number of nonzero-heritability genes (out of 17,437), defined as having (i) 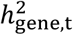 90%-CI > 0 and (ii) 90%-CI > 0 for at least one MAF bin (rare, low-frequency, or common); number (and %) of nonzero-heritability genes that explain at least 50% of cumulative 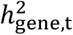 for the trait. Columns 5-7: numbers of genes with nonzero heritability contributions exclusively from common, low-frequency, or rare variants. (BMD = bone mineral density; MCH = mean corpuscular hemoglobin; MSCV = mean sphered corpuscular volume; RBC = red blood cell.)

As expected, 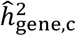 behaves similarly to 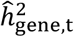. The average Pearson R^2^ of 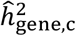 and 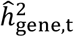 across the 25 traits is 94% (s.d. 1%) (Figure 4, Supplementary Figure 18). 92% (s.d. 1%) of nonzero-heritability genes have significant common-variant heritability; 76% (s.d. 1%) have significant causal effects exclusively from common variants (Table 1). On the other hand, 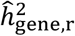 is significantly less correlated with 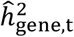 (average R^2^ = 30% (s.d. 21%) across traits) (Figure 4, Supplementary Figure 18). Approximately 2.5% (s.d. 0.6%) of genes have significant rare-variant heritability, and only 0.8% (s.d. 0.4%)—370 gene-trait pairs in total—have significant heritability exclusively from rare variants (Table 1, Supplementary Table 3). Of these 370 gene-trait pairs with only rare-variant heritability (ranging from 4 genes for heel T-score and corneal hysteresis to 32 genes for height (Table 1, Supplementary Table 3)), 232 gene-trait pairs are also identified by MAGMA^53^ (FDR < 0.05, Material and Methods). These 232 gene-trait pairs have a median 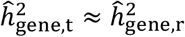 on the order of 10^-4^ whereas the median for the remaining gene-trait pairs not found by MAGMA is ~10^-6^. This suggests that MAGMA likely has limited power to detect signal from rare causal variants of moderate effect, which is expected as MAGMA tests for association between the total causal-variant signal at a gene and phenotype; it is not designed for partitioning the signal into components from different allele-frequency classes.

The 138 additional gene-trait pairs identified with our approach (Supplementary Table 4) include several genes implicated in phenotypically related Mendelian disorders. For example, *AKT2* is identified for serum gammaglutamyl transferase (90%-CI of 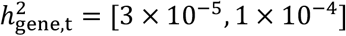, MAGMA z-score: 1.1), which is used to test for the presence of liver disease; *AKT2* is implicated in monogenic forms of type 2 diabetes^54^ and hypoinsulinemic hypoglycemia with hemihypertrophy^55^. The *AKT2* annotation used for this analysis contains a total of 104 variants; 24 are rare variants, of which 1 is identified as causal. For serum alkaline phosphatase (used to diagnose diseases related to the liver or skeletal system), we identify *MDM4* (90%-CI of 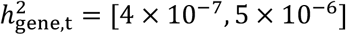, MAGMA z-score: 1.3; annotation contains 273 variants; 144 are rare variants, of which ~5 are identified as causal), which encodes a negative regulator of p53-mediated transcription^56^ that was recently implicated in an autosomal dominant bone marrow failure syndrome^57^. *COL4A4*, identified for serum apolipoprotein A1 (a test for atherosclerotic cardiovascular disease; 90%-CI of 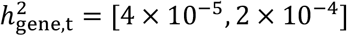; MAGMA z-score: 1.1; annotation contains 390 variants; 33 are rare variants, of which ~1 is identified as causal), is implicated in monogenic forms of kidney disease ranging in severity from hematuria to end-stage renal disease^58–61^.

We also identify several genes implicated in congenital developmental and metabolic disorders. For example, *RTTN*, identified for mean corpuscular hemoglobin (90%-CI of 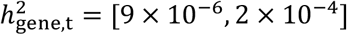; MAGMA z-score: 2.2; annotation contains 369 variants; 83 are rare, of which ~2 are identified as causal), is implicated in microcephaly, short stature, and polymicrogyria with seizures^62–65^. *SLC25A24*, identified for serum cystatin C (90%-CI of 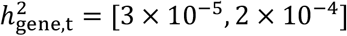; MAGMA z-score: 1.8; annotation contains 243 variants; 21 are rare, of which ~1 is causal), is implicated in Fontaine progeroid syndrome^66,67^. *TBCK*, identified for red blood cell count (90%-CI of 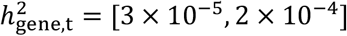; MAGMA z-score: 2.0; annotation contains 617 variants; 59 are rare, of which ~1 is causal), is implicated in infantile hypotonia with psychomotor retardation and characteristic facies^68–70^.

Taken together, these findings indicate that the rare-variant contribution to total gene-level heritability is indeed useful for identifying disease-relevant genes, especially those with moderate or relatively low total heritability, which existing methods can be underpowered to detect. Our results are consistent with the hypothesis that complex-trait variation may be explained in part by dysregulation of genes that—if completely disrupted—cause phenotypically similar or related Mendelian disorders^52^. We emphasize that, since heritability reflects genetic and phenotypic variation at the population level, if a common variant and rare variant explain the same heritability (i.e. have the same standardized causal effect size), the *allelic* effect—the expected change in phenotype per additional copy of the effect allele—is significantly larger for the rare variant.

### LoF-intolerant genes are overrepresented among genes with only rare-variant heritability

We estimate, and partition by MAF, the gene-level heritabilities of three gene sets: (i) known Mendelian-disorder genes from OMIM^49^ (n=3,446), (ii) loss-of-function (LoF)-intolerant genes (probability of LoF-intolerance (pLI) > 0.9)^50^ (n=3,230), and (iii) a set of FDA-approved drug targets for 30 immune-related traits^51^ (n=216) (Material and Methods). Compared to a set of “null” genes (sampled from the set of genes not contained in any of the three gene sets), all three gene sets have significantly higher median estimates of total and MAF-partitioned gene-level heritability (Figure 5).

**Figure 5.**
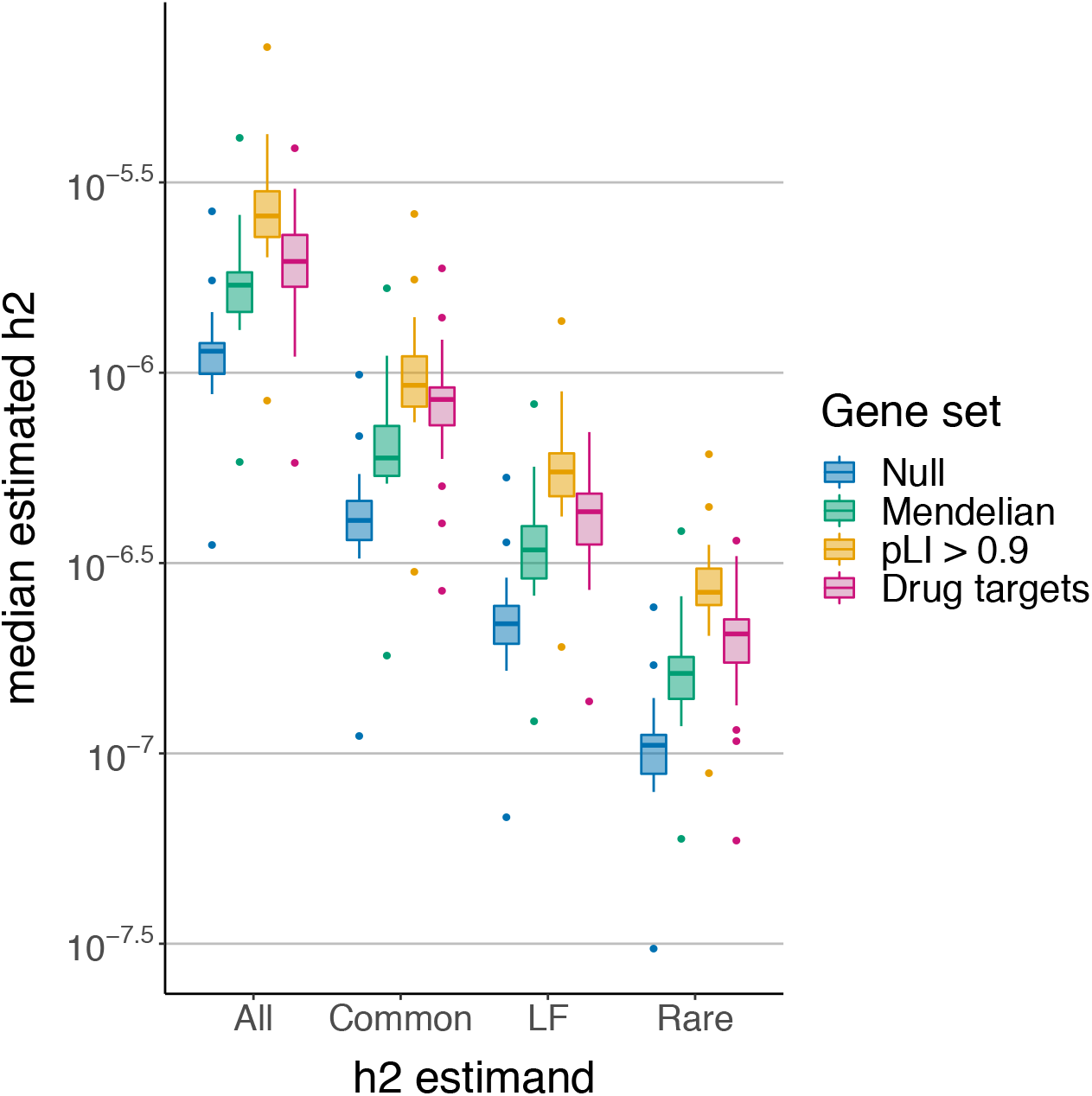
Distributions of h2 estimates for 3 gene sets. Mendelian-disorder genes (n=3,446), LoF-intolerant genes (n=3,230), and immune-related drug targets (n=216). Each point is the median posterior mean across genes for a given trait; each boxplot contains 25 quantitative traits in the UK Biobank.

We investigate whether certain classes of nonzero-heritability genes are overrepresented in the Mendelian-disorder and LoF-intolerant gene sets. The Mendelian-disorder gene set comprises ~20% of all genes and is enriched for genes with nonzero heritability for at least one trait (Fisher’s exact test, 95%-CI of OR: [1.2, 1.4]); the number of genes in both categories ranges from 261 for corneal hysteresis to 557 for height. The LoF-intolerant genes comprise ~19% of all genes and are also enriched for nonzero-heritability genes (Fisher’s exact test, 95%-CI of OR: [1.5, 1.7]); the overlap between the two categories ranges from 314 genes for corneal hysteresis to 650 for height. In contrast, genes with exclusively rare-variant heritability are significantly enriched in the LoF-intolerant gene set (95%-CI of OR: [1.1, 2.1]) but not in the Mendelian-disorder gene set (95% CI of OR: [0.9, 1.7]). On average across traits, ~19% (s.d. 11%) of the previously identified 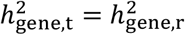 genes and ~21% (s.d. 1%) of genes with only common-variant heritability are also in the Mendelian-disorder gene set. In contrast, ~32% (s.d. 16%) of genes with 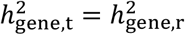 are also in the LoF-intolerant gene set, compared with ~23% (s.d. 1%) of genes with 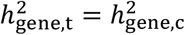.

### MAF-partitioned gene-level heritability reveals unique insights into genetic architecture

We investigated whether gene-level heritability estimates are correlated with gene length, average LD score of variants in the gene (a proxy for the strength of LD in the region), and average MAF of variants in the gene. 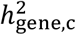 (and, to a large extent, 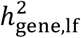) is distributed very similarly to 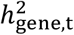 with respect to these variables (Figure 6, Supplementary Figure 19). However, the distribution of 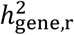 shows marked differences, particularly with respect to gene length. Specifically, we observe higher average 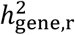 among shorter genes even though the number of causal variants per gene (across all allele frequencies) increases with gene length (Figure 6, Supplementary Figure 20). The expected per-causal variant effect size per gene is invariant to gene length for common and low-frequency variants, but for rare variants, the average across gene-trait pairs is nearly 10^-4^ in the shortest quintile of genes versus 10^-6^ in the longest (Figure 6). While this result initially seems paradoxical, it is not inconsistent with the literature; previous studies have reported strong inverse correlations between gene length and expression which could be due to, for example, natural selection favoring fewer/shorter introns in highly expressed genes due to the high energy/costs associated with transcription and splicing^71,72^.

**Figure 6.**
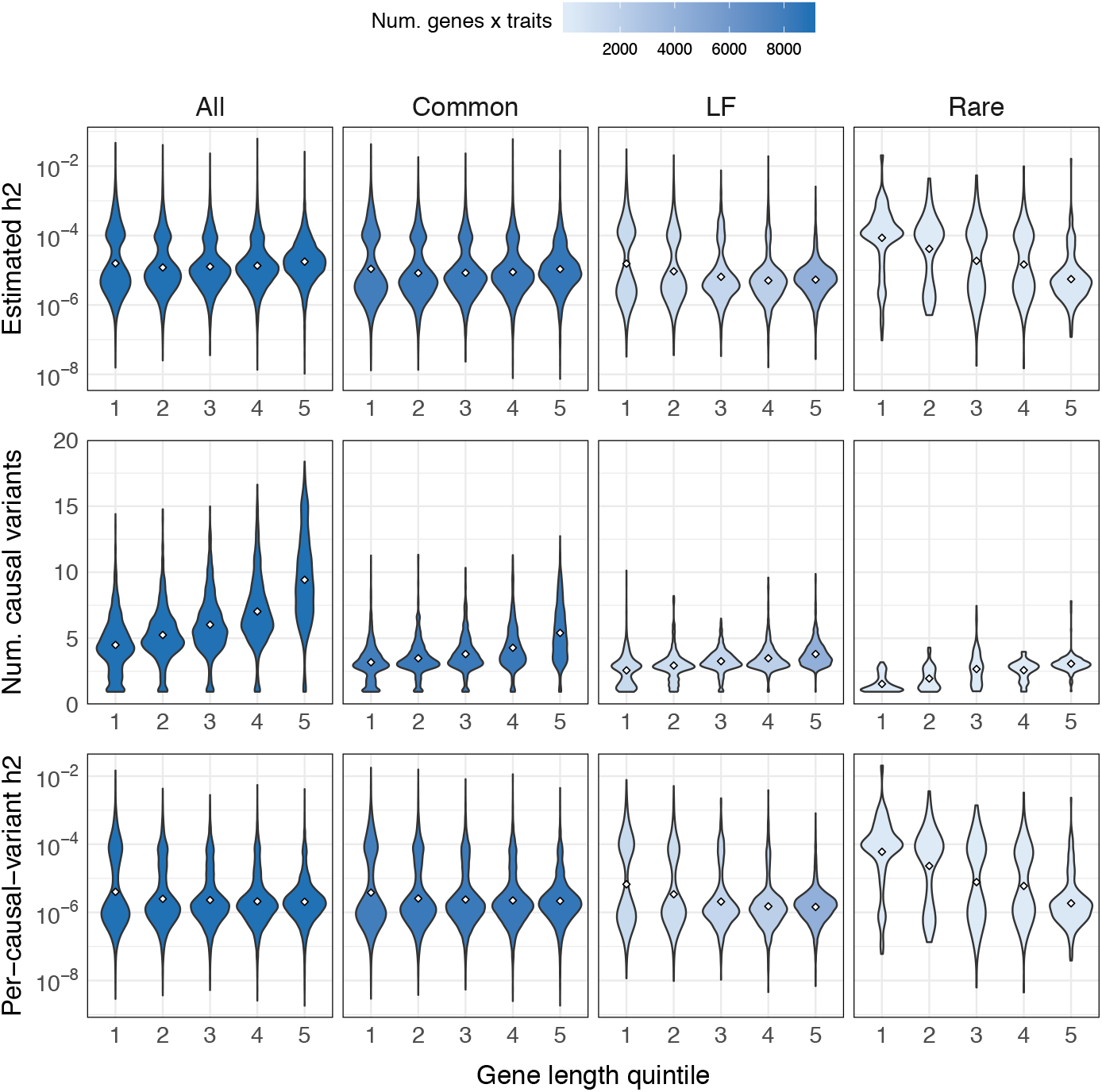
Inverse relationship between rare-variant h2 estimates and gene length. Estimates of h2 (top), number of causal variants per gene (middle), and expected effect size per causal variant per gene (bottom) with respect to gene length (x-axis) for 25 traits. Each violin plot is the distribution of posterior mean estimates for nonzero-heritability genes with 90%-CIs > 0 for each h2 quantity. Color gradient indicates the number of estimates in each violin plot (number of gene-trait pairs).

Using the empirical distributions of cumulative 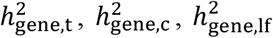, and 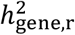, we loosely quantify differences in polygenicity at the level of genes (with the caveat that, since there is a high degree of gene overlap in some regions, cumulative 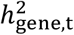 may be more informative for some traits over others) (Figure 7). For example, if cumulative 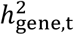 is divided equally among nonzero-heritability genes, the empirical CDF for 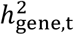 would be the line *y* = *x*, where the x-axis is the rank ordering of genes from highest to lowest 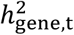; two traits with the same empirical CDF for 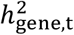 can have different empirical CDFs for each MAF-partitioned component. Once again, we find that the cumulative distributions of 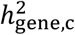 are extremely similar to those of 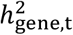 (Figure 7, Supplementary Figure 21). Although the curves generally have similar shapes across traits (i.e. similar spread of heritability across genes), some traits have a notable amount of heritability concentrated in just the top gene, and many of these gene-trait pairs have been functionally validated in the literature. For example, for serum urate concentration, *SLC2A9* — a known urate transporter^73–75^ — is the single largest contributor to total, common-, and LF-variant gene-level heritability 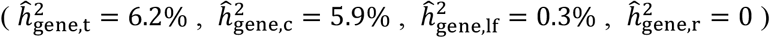, accounting for 46%, 51%, and 29% of the cumulative heritability for each estimand, respectively (Figure 7); certain loss-of-function mutations in *SLC2A9* are known to cause a rare form of renal hypouricemia^76–78^, a disorder characterized in part by low serum urate levels. For serum alkaline phosphatase, we find that *ALPL* — which encodes the enzyme alkaline phosphatase — is the single largest contributor to total and LF-variant gene-level heritability 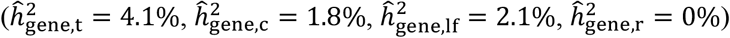, explaining 15% and 39% of the respective cumulative heritability estimands (Figure 7); certain loss-of-function mutations in *ALPL* are known to cause hypophosphatasia, a monogenic disorder characterized in part by low alkaline phosphatase^79,80^.

**Figure 7.**
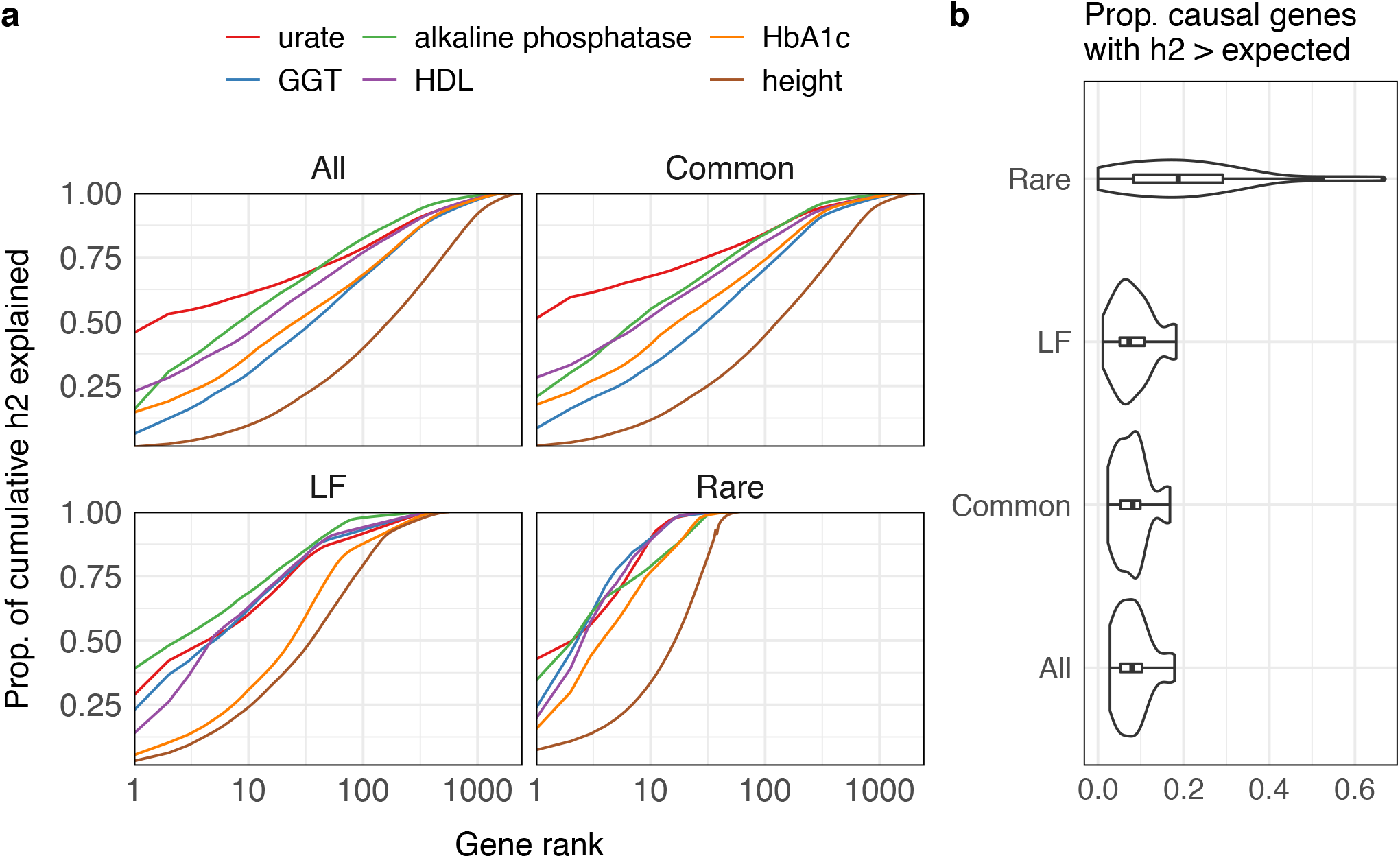
Gene-level heritability estimates capture differences polygenicity across traits. (a) Empirical distributions of cumulative heritability for six example traits (clockwise from top left: total, common, low-frequency, and rare). Each curve can be read as, “the top X genes explain Y% of the cumulative gene-level heritability for a given trait.” Cumulative gene-level h2 is estimated by summing the estimated posterior means for nonzero-h2 genes (90%-CI > 0). (Supplementary Figure 21 shows all 25 traits.) (b) Proportion of nonzero-h2 genes per trait with disproportionately large heritability estimates, defined as genes with 90%-CI > (cumulative heritability / number of causal genes)). Each violin plot represents 25 traits.

## Discussion

We propose a general approach for estimating the heritability explained by any set of variants much smaller than an LD block and assess its utility in estimating/partitioning gene-level heritability. In simulations, we confirm that incorporating uncertainty about which variants are causal and what their effect sizes are dramatically improves specificity over naive approaches that ignore uncertainty in the causal effects. For 25 complex traits and >17K genes, we estimate gene-level heritability—the heritability explained by variants in the gene body plus a 10-kb window upstream/downstream from the gene start/end positions—and partition by allele-frequency class to explore differences in genetic architecture across traits. As expected, most gene-level heritability is dominated by common variants, but we identify several genes with nonzero heritability exclusively from rare or low-frequency variants. Notably, we identify many genes with nonzero gene-level heritability explained exclusively by rare variants that existing methods are underpowered to detect. Many of these genes have known roles in Mendelian disorders that are phenotypically similar or related to the complex trait; we also identify genes implicated in systemic congenital developmental and metabolic disorders. Our results demonstrate that the rare-variant contribution to total gene-level heritability is a useful quantity that can be considered alongside common-variant heritability enrichments to obtain a more comprehensive understanding of genetic architecture.

We conclude by discussing the limitations of our approach. First, multiple lines of evidence suggest that rare and “ultra-rare” variants, which are not well-tagged by variants on genotyping arrays, may explain much of the “missing heritability” not captured by genotyped or imputed variants^81–84^. Since imputed genotypes are noisier for rarer variants and variants in lower LD regions, we analyze variants with MAF > 0.5%. Additional work is needed to assess the error incurred by using genotyped/imputed data in lieu of whole genome sequencing (WGS) as well as the signal that is missed by excluding variants with MAF < 0.5%. While our estimator can be applied to whole exome sequencing (WES) data, LD between coding and noncoding regions would significantly inflate gene-level heritability estimates; LD between exonic and intronic variants could also cloud interpretation, depending on the application. With multiple biobanks starting to sequence large numbers of individuals^85–88^, we believe the availability of large-scale WGS data will gradually become less of an issue.

We correct for population structure using genome-wide principal components (PCs) computed from the same imputed genotypes that are used to perform each GWAS. This is a standard approach to correcting for population stratification, which typically reflects geographic separation, in estimates of genome-wide SNP-heritability and genome-wide functional enrichments, both of which are driven by common SNPs. However, rare variants generally have more complex spatial distributions and thus exhibit stratification patterns distinct from those of common SNPs^84,89^. It is unclear whether methods that are effective for controlling stratification of common SNPs are applicable to rare variants^90^. We leave the question of whether uncorrected structure among rare variants significantly influences our estimates of gene-level heritability for future work.

Our approach requires OLS association statistics and LD computed from a subset of individuals in the GWAS. While estimates of gene-level heritability and the MAF-partitioned components are robust to sample sizes as low as 5,000, the individuals used to estimate LD must be a subset of the individuals in the GWAS. Although summary association statistics are publicly available for hundreds of large-scale GWAS, most of these studies are meta-analyses and therefore do not have in-sample LD available. Moreover, many publicly available summary statistics were computed from linear mixed models rather than OLS, which is used throughout our simulations and derivations. Additional work is needed to extend our approach to allow external reference panel LD (e.g., 1000 Genomes^91^) and/or mixed model association statistics. Biobanks can help to ameliorate potential issues stemming from noisy LD by releasing summary LD information in addition to summary association statistics^92^.

Finally, gene-level heritabilities of different genes can have nonzero covariance due to physical overlap between genes and/or correlated causal effect sizes. Thus, the heritability estimates reported in this work have additional sources of noise/uncertainty which were not directly modeled or accounted for. Since modeling correlation of causal effect sizes would make inference considerably more challenging, we leave this for future work. Importantly, genes with credible intervals > 0 should not be interpreted as “causal” for the complex trait without additional functional validation, as nonzero gene-level heritability indicates association but not causality.

## Acknowledgments

We thank the UK Biobank Resource (application #33297) for making this work possible. We are also grateful to Alkes Price, Harold Pimentel, Luke O’Connor, Nasa Sinnott-Armstrong, and Ruth Johnson for providing helpful comments and discussion. This work was funded in part by the National Institutes of Health under awards R01-HG009120, R01-MH115676.

## Web Resources

Protein-coding gene list: https://github.com/bogdanlab/gene_sets/protein_coding_genes.bed

OMIM gene list: https://github.com/bogdanlab/gene_sets/blob/master/mendelian_genes.bed

LoF-intolerance metrics by gene: https://gnomad.broadinstitute.org/downloads

susieR software: https://github.com/stephenslab/susieR

MAGMA software: https://ctg.cncr.nl/software/magma

PLINK software: https://www.cog-genomics.org/plink2

The UK Biobank Resource: https://www.ukbiobank.ac.uk/

## Material and Methods

### Model and definitions of estimands

We model the phenotype of a given individual using a standard linear model, *y* = **x**^T^**β** + *ϵ*, where **x**^T^ = (*x* … *x_M_*)^T^ is the vector of standardized genotypes at M variants, i.e. 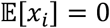 and *var*[*x_i_*] = 1 for *i* = 1, …, *M*. **β** is the M × 1 vector of standardized causal effect sizes, and 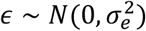 is environmental noise. We assume that the phenotype is standardized in the population, i.e. 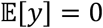, *var*[*y*] = 1. Linkage disequilibrium (LD) between variants *i* and *j* is defined as *r_i, j_* ≡ *cov*[*x_i_ x*_j_] = 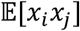 and the full LD matrix for all M variants is **R** ≡ *cov*[**x**^T^].

Letting *p*_causal_ ∈ [0, 1] such that *M* × *p*_causal_ is the total number of causal variants, we assume the causal effect of the *i*-th variant is 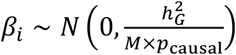 with probability *p*_causal_ or *β_i_* = 0 with probability 1 – *p*_causal_. Under this model, total SNP-heritability 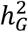 is defined as the proportion of phenotypic variance explained by the M variants,

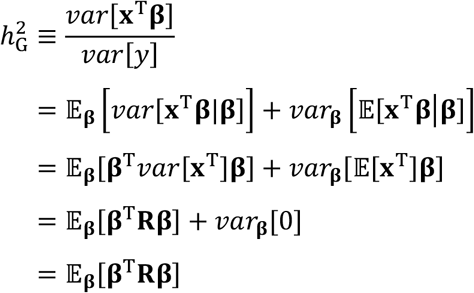

where the second line follows from the Law of Total Variance.

Let *g* index a gene of interest. Given an assignment of *m_g_* variants to gene *g*, let 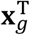 be the *m_g_* × 1 vector of genotypes at this set of variants and let 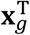, be the genotypes of the remaining *M* – *m_g_* variants. We can rewrite the total SNP-heritability of the trait in terms of gene *g* as

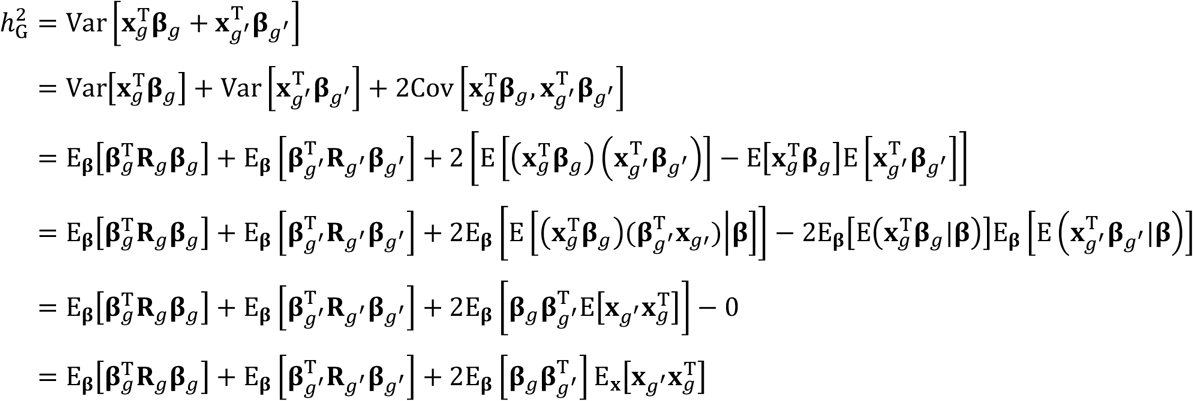

where the fourth line follows from the Law of Total Expectation. If we additionally assume that *cov*[*β_i_*, *β_j_*] = 0 for all *i* ≠ *j*, then 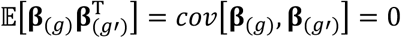, which simplifies the above equation to

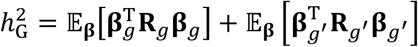

We refer to the first term, the component of heritability attributable to the causal effects in gene *g*, as *total genelevel heritability*, i.e.

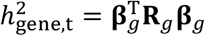

Using the same assumptions as above, we can partition the variants in gene *g* by minor allele frequency such that

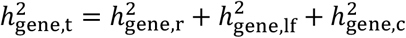

where 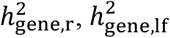, and 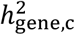 are the components of 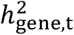 attributable to the causal effects of rare (MAF < 0.01), low-frequency (0.01 ≤ MAF < 0.05), and common (MAF ≥ 0.05) variants, respectively. The estimands of interest in this work are the four terms in 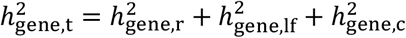.

### Estimating the posterior distribution of gene-level heritability

Since we have neither the “true” causal effect sizes, **β**, nor the population LD, **R**, we must estimate both from data. We consider one approximately independent LD block at a time. Given a GWAS of *N* individuals, let 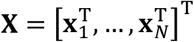 be the *N* × *M* matrix of standardized genotypes measured at *M* variants, let **y** = (*y*_1_, …, *y_N_*)^T^ be an *N* × 1 vector of phenotypes, and let 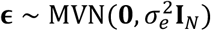 be environmental noise.

It is often the case that individual-level genotype data are inaccessible for privacy or logistical reasons. However, GWAS summary statistics—estimates of the causal effects and their standard errors—are publicly available for thousands of traits. Ordinary least squares (OLS) estimates of the causal effects are often provided, defined as

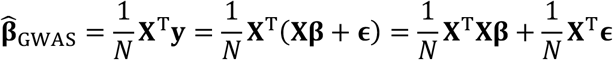

It follows that

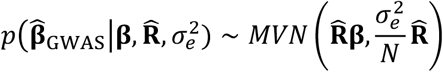

In this scenario, the observed data D are not the individual-level genotypes and phenotypes (**X, y**), but rather 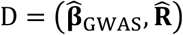, where 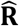 is an estimate of LD computed from either the genotypes of a set of individuals in the GWAS (“in-sample” LD) or from an external reference panel (e.g., 1000 Genomes^91^). By combining the prior on **β**, *p*(**β**|***λ***) (***λ*** represents hyperparameters in the prior over **β**, estimated with empirical Bayes procedure as implemented in SuSiE), and the likelihood of the observed data, 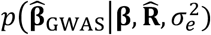, one can compute the posterior distribution of the causal effects, 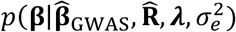. The hyperparameters ***λ*** and 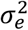 can be estimated with an empirical Bayes procedure as in SuSiE framework. We note that for computational efficiency, we can partition the whole genome into approximately independent LD blocks, and estimate the posterior distribution of **β** separately for each LD block. Because each LD block is approximately independent of the rest of the genomes by definition, the genetic effects from SNPs outside of the LD block of interest are absorbed into the environmental noise. And correspondingly, the LD block-specific hyperparameters (***λ***, 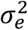) are estimated independently for each LD block.

The posterior of **β**, *p*(**β**|D), is in general computationally intractable. Approximate inference, e.g., Markov Chain Monte Carlo (MCMC) or variance inference, can be used to approximate the exact posterior *p*(**β**|D) as 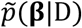. In this work, we use SuSiE^47^, a variational inference-based implementation of linear regression with sparse prior. (In principle, it is straightforward to use other implementations of linear regression with sparse prior). We draw *K* samples from the posterior of the causal effects, 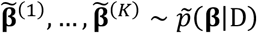. This approximate distribution can in turn be used to approximate the full posterior distribution of 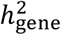, i.e. 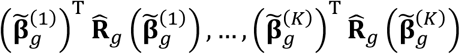.

Finally, given the approximate posterior of 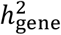, one can compute the posterior mean,

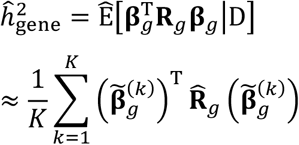

and measures of uncertainty such as credible intervals (described below). Similar procedures could be applied to estimate the gene-level heritabilities stratified by annotations of SNPs (such as MAF-based annotation).

### Quantifying uncertainty in gene-level heritability estimates

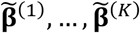 provide an approximation to the full posterior distribution of **β**, thus capturing *uncertainty* about the causal effect sizes arising from two main sources: LD and finite GWAS sample size (Figure 1). Therefore, by using the full posterior of **β** to approximate the full posterior of 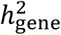, we wish to capture uncertainty in the causal effects that propagates into our estimate of 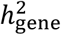. (The noise in 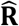 is also an important factor but, for simplicity, we first investigate uncertainty in 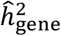 in simulations where 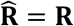.)

We summarize the uncertainty in 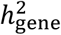 by computing *ρ*-level credible intervals (*ρ*-CIs). For a given *ρ* ∈ [0, 1], *ρ*-CI is defined as the central interval within which 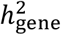 lies with probability *ρ*, i.e. the upper and lower bounds of *ρ*-CI are set to the empirical 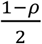 and 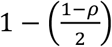 quantiles of the posterior samples 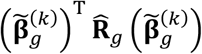, *k* = 1, …, *K*.

### Implementation details

We partition the genome into approximately independent LD blocks^93^ and, for each gene of interest, we perform inference on the LD block containing the gene. For each LD block, we extract the marginal association statistics and estimate LD for all the variants in the LD block. We estimate the posterior distribution of effect sizes using the function “susie_suff_stat” with default parameters, as implemented in SuSiE^47^ v0.8. We use the function “susie_get_posterior_samples” to obtain 500 posterior samples.

### Simulation framework

We obtain the real imputed genotypes of N=290,273 “unrelated white British” individuals in the UK Biobank by extracting individuals with self-reported British ancestry who are > third-degree relatives (pairs of individuals with kinship coefficient < ½^(9/2)^, as defined in ref.^48^). Filtering on MAF > 0.5% leaves 200,235 variants on chromosome 1. A list of 1,083 genes on chromosome 1 and their coordinates were downloaded from https://github.com/bogdanlab/gene_sets (Data Availability). For each variant, genotypes are standardized such that the mean is 0 and variance is 1 across individuals. Phenotypes were simulated under a variety of genetic architectures according to the following steps. First, we randomly select 3%, 8%, or 16% (out of the 1,083 genes) to be causal 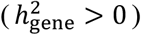. Second, we draw causal variants in the causal gene bodies and within 10-kb upstream/downstream of the gene start/end positions; the causal variants in the window around the gene are intended to represent regulatory causal variants in transcription start sites (TSSs). The causal configuration is set to be either (1) 5 causal variants in gene body and 3 causal variants in TSS or (2) 10 causal variants in gene body and 6 causal variants in TSS. Third, we draw noncoding “background” causal variants across the whole chromosome with frequency *p*_causal_ = {0.001, 0.01}. Finally, conditional on the causal statuses of the variants, we draw independent causal effect sizes from a Gaussian distribution where the variance of each causal variant is standardized such that the gene bodies collectively have a heritability of 3%, TSSs collectively have 1%, and noncoding background variants together explain 1%. We note that the causal statuses and effect sizes for each variant are only drawn once; the environmental noise term is drawn 30 times independently to generate 30 simulation replicates.

### Evaluating and comparing gene-level heritability estimates in simulations

Recall that for a given gene *g*, the causal effect sizes and LD of the variants assigned to the gene are denoted **β**_*g*_ and **R**_*g*_, and ground-truth gene-level heritability is defined as 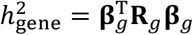. The posterior mean estimated for a single simulation replicate *s* is denoted 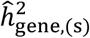. We estimate the bias of the estimator as 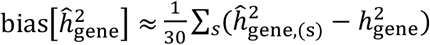; the variance of the estimator as 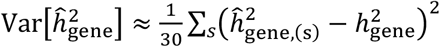; and the mean 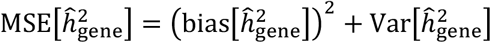.

For each simulation replicate *s*, we also output *ρ*-level credible intervals, defined as 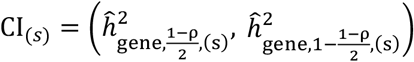 where the 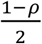 and 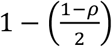 quantiles are estimated from the posterior samples. To assess the accuracy of credible intervals, we calculate *empirical coverage* across simulation replicates, defined as the proportion of simulation replicates in which the *ρ*-level credible interval covers the ground-truth gene-level heritability: 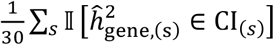.

### Comparison to “naïve” gene-level heritability estimator

We compare our approach to an alternative “naïve” estimator of gene-level heritability that does not model LD between the gene and its adjacent regions and thus ignores causal-effect uncertainty. This estimator is similar to existing methods that are meant to be applied to approximately independent LD blocks^45,94^. For each gene *g*, we extract the marginal association statistics, 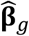, and the estimated LD, 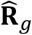, for the variants assigned to the gene, and we compute the alternative estimator as 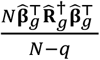, where 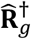 and *q* are the pseudo-inverse and rank of 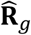, respectively^45,94^.

### Assessing robustness to LD panel sample size

To assess the robustness of our approach to the sample size of the LD panel used to estimate LD, we randomly draw a subset of N={500, 1000, 2500, 5000} individuals from the full 290,273 individuals. After extracting variants with MAF > 0.5%, genotypes are standardized to have mean 0 and variance 1, similar to the full-sample analysis. Since we are interested in assessing robustness to noisy estimates of LD, all analyses are performed using the same set of marginal association statistics used in the full-sample analysis, excluding the variants that were filtered from the LD panel based on MAF. The LD and marginal association statistics are fed into the *h2gene* software, similar to the full-sample analysis.

### Analysis of 25 UK Biobank phenotypes

We analyzed 25 quantitative phenotypes in the UK Biobank. Phenotypes and imputed genotypes were filtered according to the same procedures used in the simulation analyses, leaving N=290,273 individuals and M=5,650,812 variants with MAF > 0.5%. Quantitative phenotypes were quantile-normalized to a Gaussian distribution with mean 0 and variance 1. We then performed a GWAS for each trait using the “assoc” option in PLINK (Web Resources) with age, sex, and the top 10 genetic principal components included as covariates. We computed in-sample LD for each approximately independent LD block^93^. We downloaded gene names and coordinates from https://github.com/bogdanlab/gene_sets and, for each gene, we define the estimand of interest to be a function of the variants in the gene body *and* those located within 10-kb upstream/downstream of the gene start/end positions. Finally, given the in-sample LD and marginal association statistics, we infer the posterior distribution of the causal effect sizes one LD block at a time, and we estimate and partition gene-level heritability for all genes in each LD block, where we define the estimand of interest to be a function of the variants in the gene body *and* those located within 10-kb upstream/downstream of the gene start/end positions. MAGMA v1.09 was used for gene-level association with a 10kb window around each gene. The same list of genes and the same set of imputed variants were used for the MAGMA analysis.

