## Supplemental Information for "Partitioning gene-level contributions to complex-trait heritability by allele frequency identifies disease-relevant genes"

August 17, 2021

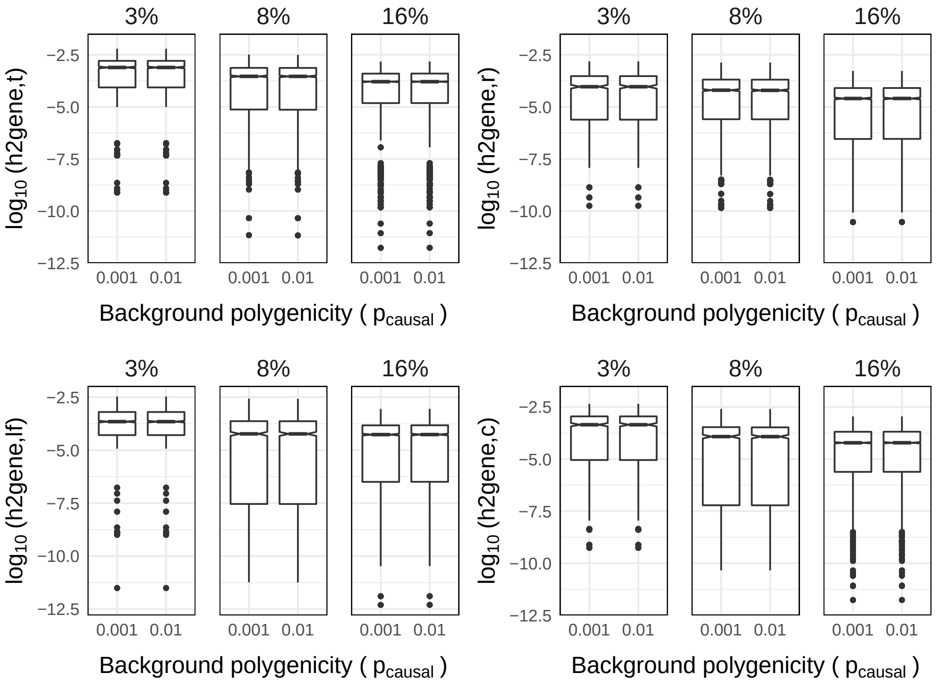


**Supplementary Figure 1**: Distributions of true $h^2_{\text{gene,t}}$ estimands for causal genes in simulations on chromosome 1. Total $h^2_G = 0.05$. Cumulative $h^2_{\text{gene,t}} = 0.03$. 30 simulation replicates, MAF $>$ 0.005, N=291K. The proportion of causal genes was 3%, 8%, or 16% drawn from a total of 1,083 genes.

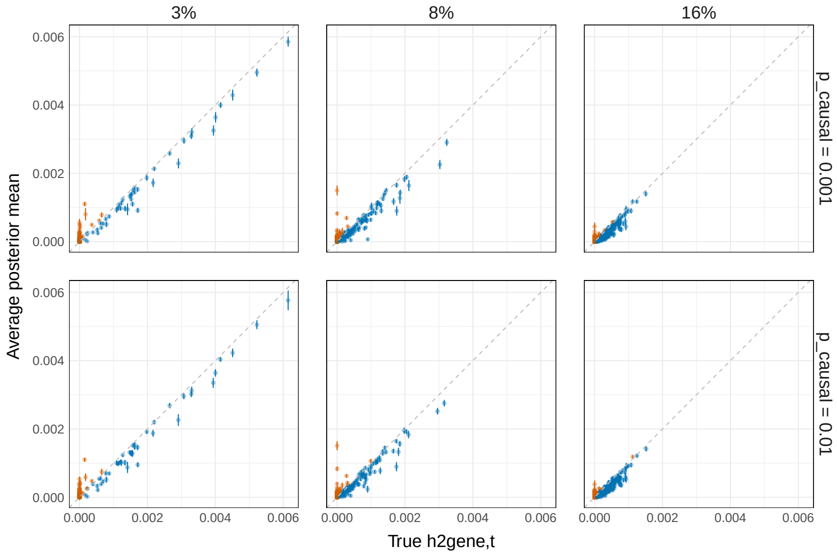


**Supplementary Figure 2**: Average $\hat{h}^2_{gene}$ vs. true $h^2_{gene}$ in simulations on chromosome 1 where 3%, 8%, or 16% of genes are causal. Orange points are genes with significantly upward-biased $\hat{h}^2_{gene}$; blue points are all other genes. Bias is estimated as the average error, $(\hat{h}^2_{gene} - h^2_{gene})$, from 30 simulation replicates, and is considered "significant" if the error bars ($\pm 1.96$ s.e.m.) do not overlap the true value of $h^2_{\text{gene,t}}$. Total $h^2_G = 0.05$. Cumulative $h^2_{gene} = 0.03$. 30 simulation replicates, MAF $>$ 0.005, N=291K.

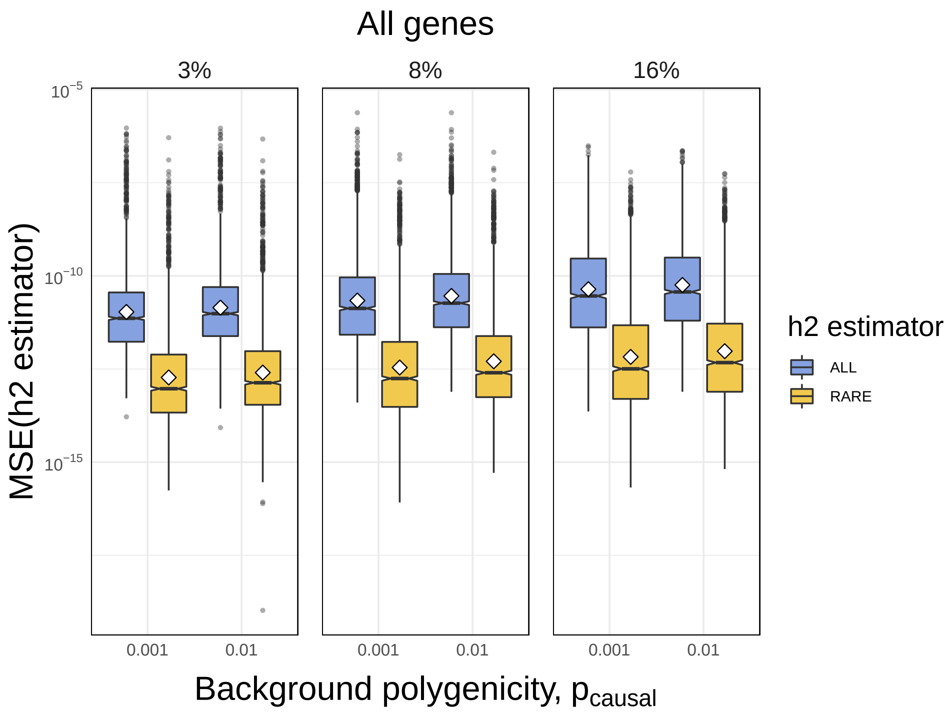


**Supplementary Figure 3**: MSE of $\hat{h}^2_{\mathrm{gene,t}}$ (blue) and $\hat{h}^2_{\mathrm{gene,r}}$ (yellow) for all 1,083 genes with respect to background polygenicity (x-axis) in simulations on chromosome 1 where either 3%, 8%, or 16% of genes are causal. Each point in each boxplot is the MSE for a single gene estimated from 30 simulation replicates. Total $h^2_G = 0.05$. Cumulative $h^2_{\mathrm{gene,t}} = 0.03$, MAF $>$ 0.005, $N = 291K$.

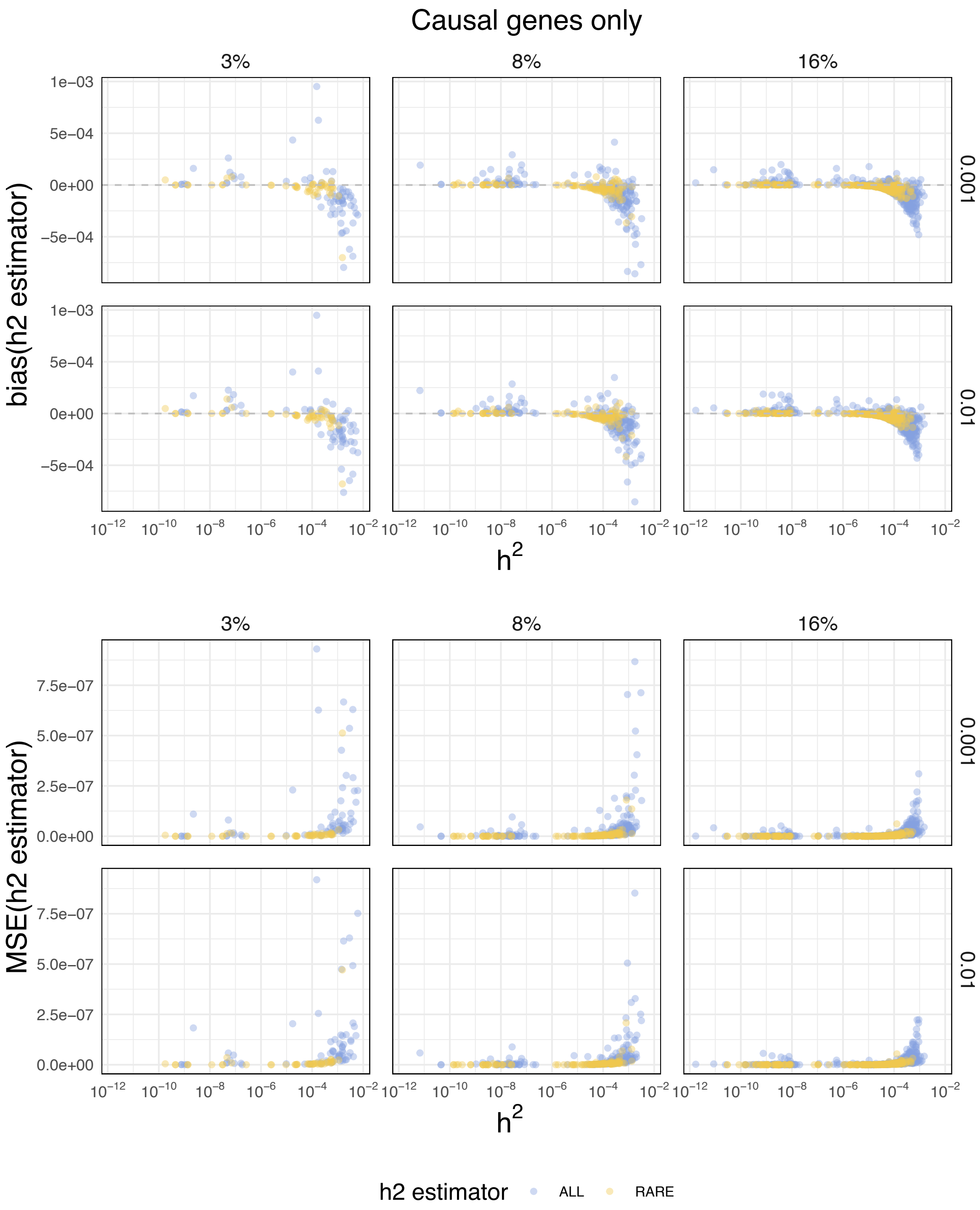


**Supplementary Figure 4**: Bias (top) and MSE (bottom) with respect to true $h^2$ (x-axes) in simulations on chromosome 1 where either 3%, 8%, or 16% of genes are causal. Only causal genes are shown ($h^2_{\text{gene,t}} > 0$ for blue; $h^2_{\text{gene,r}} > 0$ for yellow). Total $h^2_G = 0.05$. Cumulative $h^2_{gene} = 0.03$, MAF > 0.005, $N = 291K$.

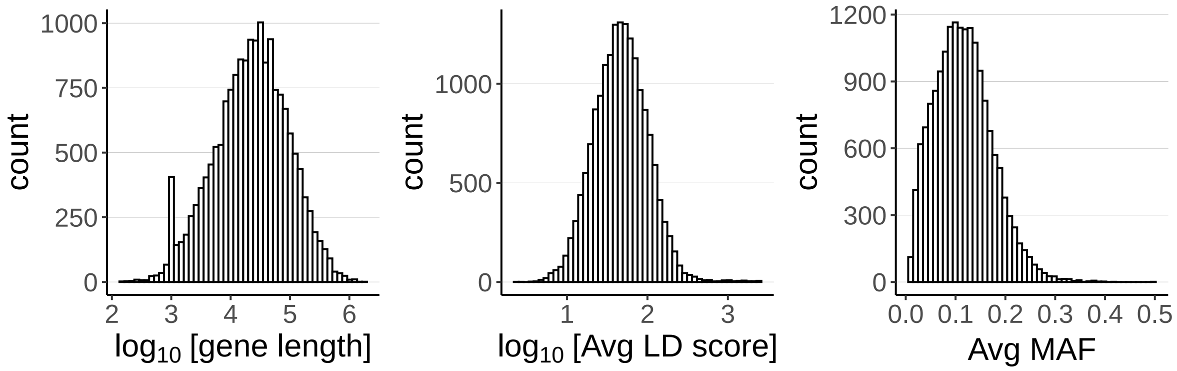


**Supplementary Figure 5**: Distribution of gene lengths (left), average LD score of variants assigned to gene (middle), and average MAF of variants assigned to gene (right) for 17,437 genes.

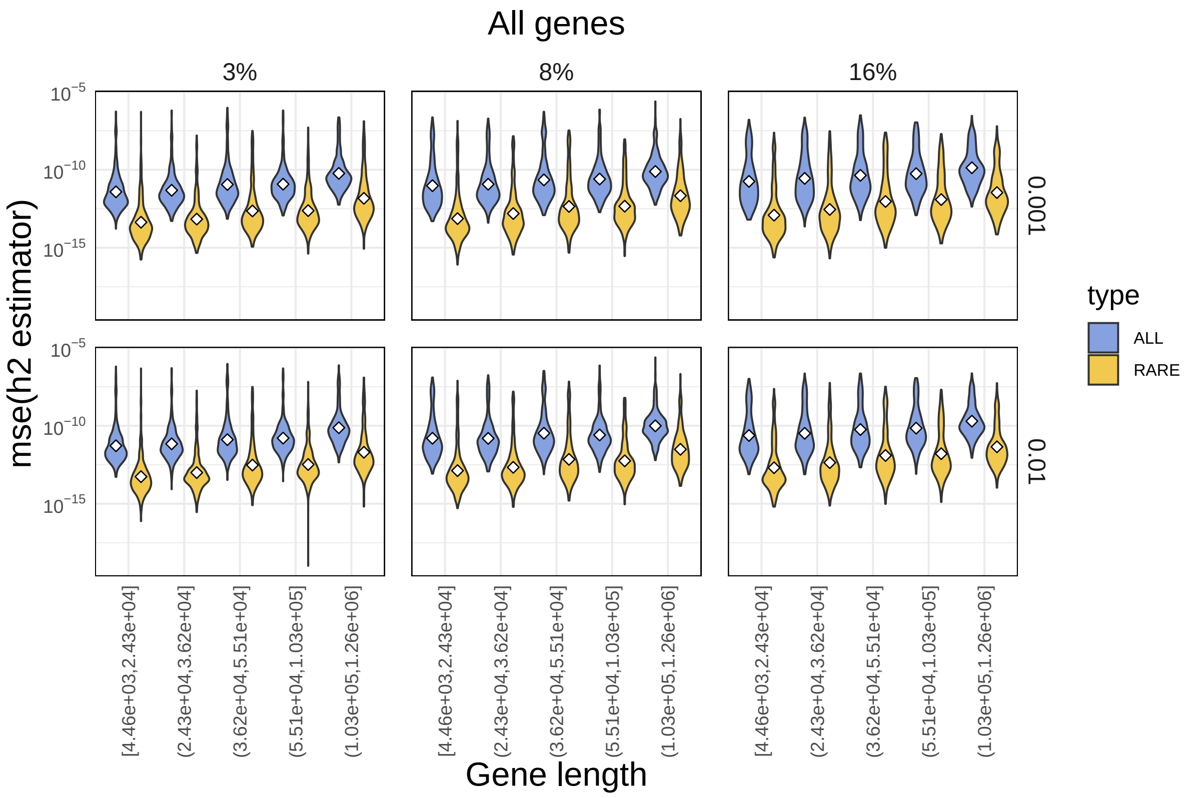


**Supplementary Figure 6**: MSE with respect to gene length. Total $h^2_G = 0.05$. Cumulative $h^2_{\text{gene,t}} = 0.03$, MAF > 0.005, $N = 291K$.

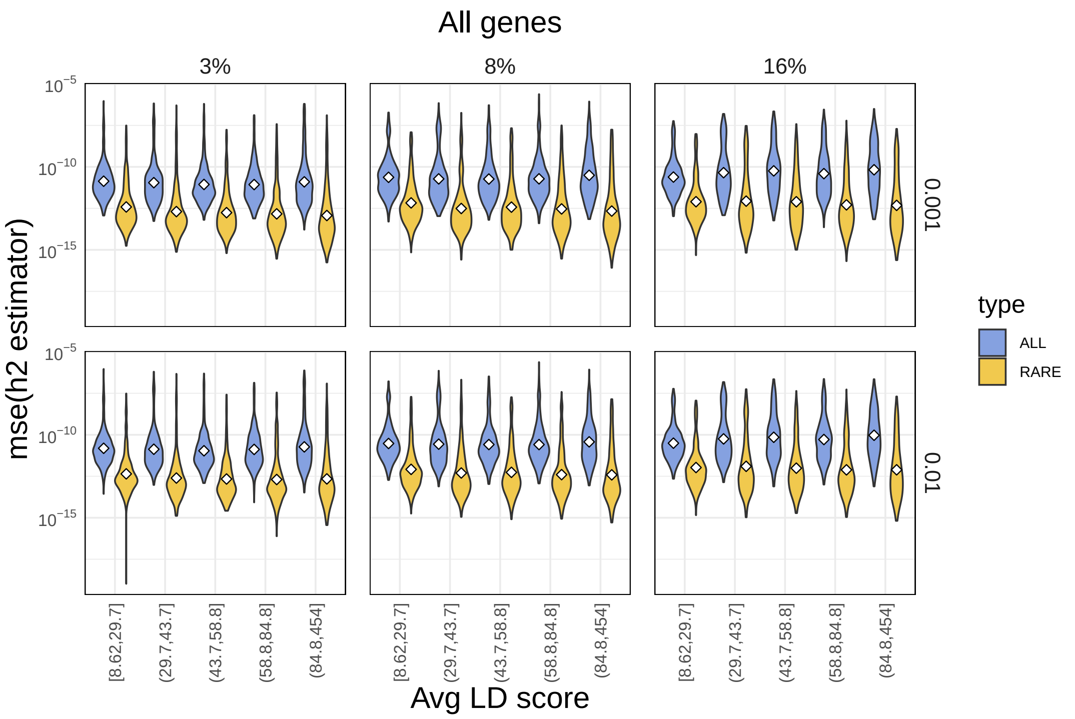


**Supplementary Figure 7**: MSE with respect to average LD score of variants assigned to gene. Total $h_G^2 = 0.05$. Cumulative $h^2_{\text{gene,t}} = 0.03$, MAF > 0.005, $N = 291K$.

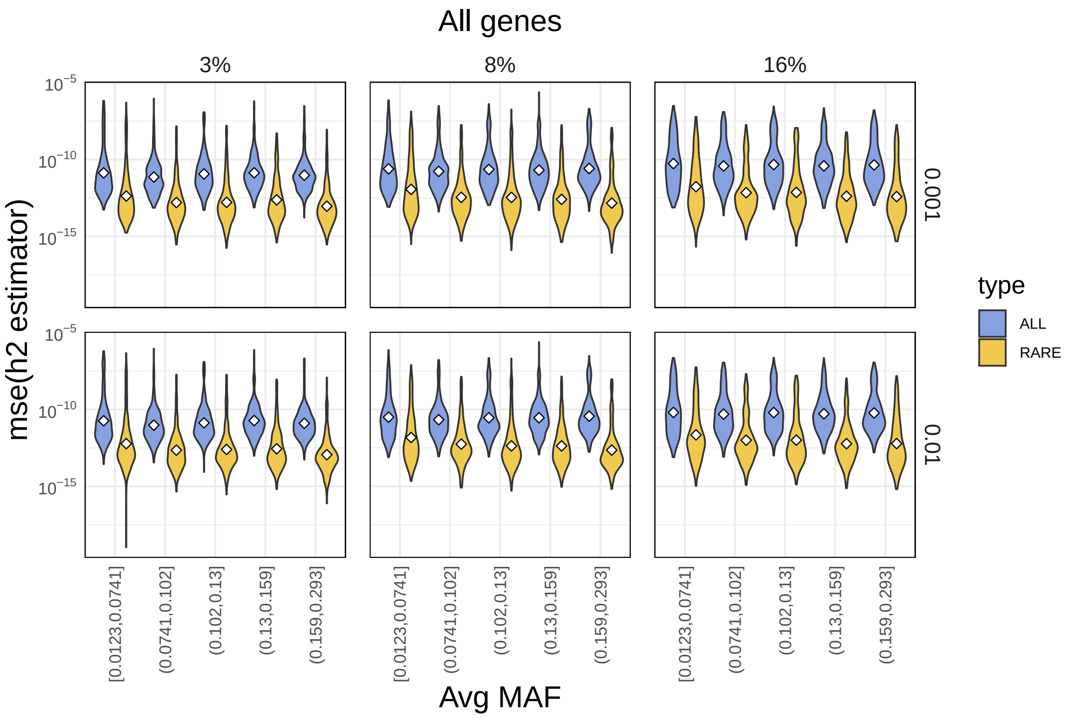


**Supplementary Figure 8**: MSE with respect to average MAF of variants assigned to gene. Total $h_G^2 = 0.05$. Cumulative $h^2_{\text{gene,t}} = 0.03$, MAF > 0.005, $N = 291K$.

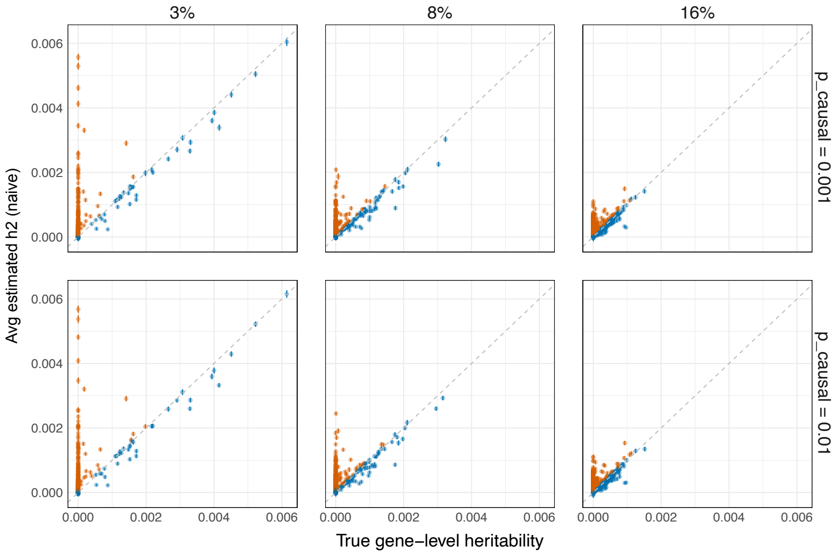


**Supplementary Figure 9**: Average estimated $h^2_{\mathrm{gene,t}}$ from "naive method" vs. true $h^2_{\mathrm{gene,t}}$ in simulations on chromosome 1 where 3%, 8%, or 16% of genes are causal. Orange points are genes with significant upward bias; blue points are all other genes. Bias is estimated as the average error from 30 simulation replicates and is considered "significant" if the error bars ($\pm 1.96$ s.e.m.) lie above the true value of $h^2_{\mathrm{gene,t}}$. Total $h^2_G = 0.05$, cumulative $h^2_{\mathrm{gene,t}} = 0.03$, MAF $>$ 0.005, N=291K.

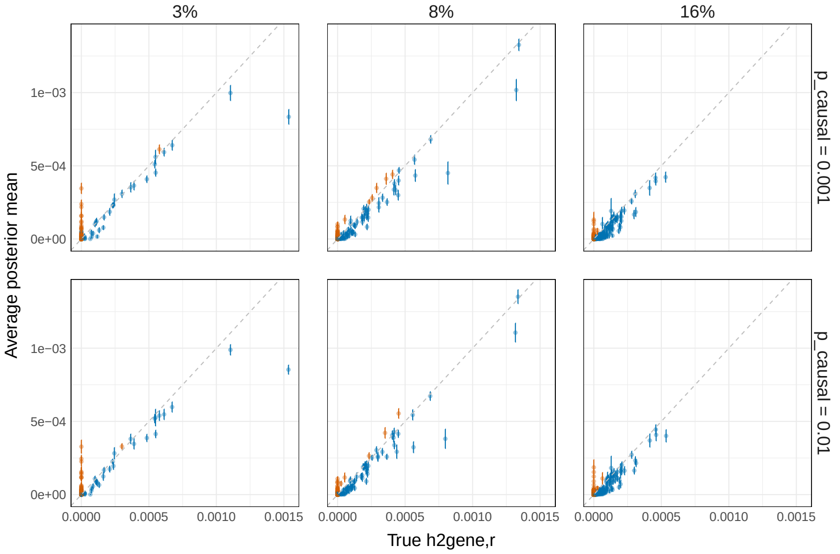


**Supplementary Figure 10**: Average estimated posterior mean of $h^2_{\mathrm{gene,r}}$ vs. true $h^2_{\mathrm{gene,r}}$ in simulations on chromosome 1 where 3%, 8%, or 16% of genes are causal. Orange points are genes with significant upward bias; blue points are all other genes. Bias is estimated as the average error from 30 simulation replicates and is considered "significant" if the error bars ($\pm 1.96$ s.e.m.) lie above the true value of $h^2_{\mathrm{gene,r}}$. Total $h^2_G = 0.05$, cumulative $h^2_{\mathrm{gene,t}} = 0.03$, MAF $>$ 0.005, N=291K.

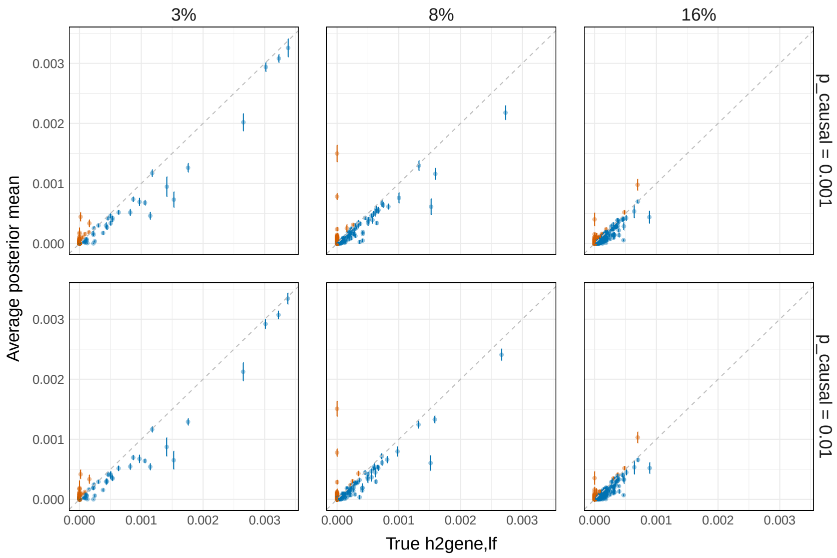


**Supplementary Figure 11**: Average estimated posterior mean of $h^2_{\mathrm{gene,lf}}$ vs. true $h^2_{\mathrm{gene,lf}}$ in simulations on chromosome 1 where 3%, 8%, or 16% of genes are causal. Orange points are genes with significant upward bias; blue points are all other genes. Bias is estimated as the average error from 30 simulation replicates and is considered "significant" if the error bars ($\pm 1.96$ s.e.m.) lie above the true value of $h^2_{\mathrm{gene,lf}}$. Total $h^2_G = 0.05$, cumulative $h^2_{\mathrm{gene,t}} = 0.03$, MAF $>$ 0.005, N=291K.

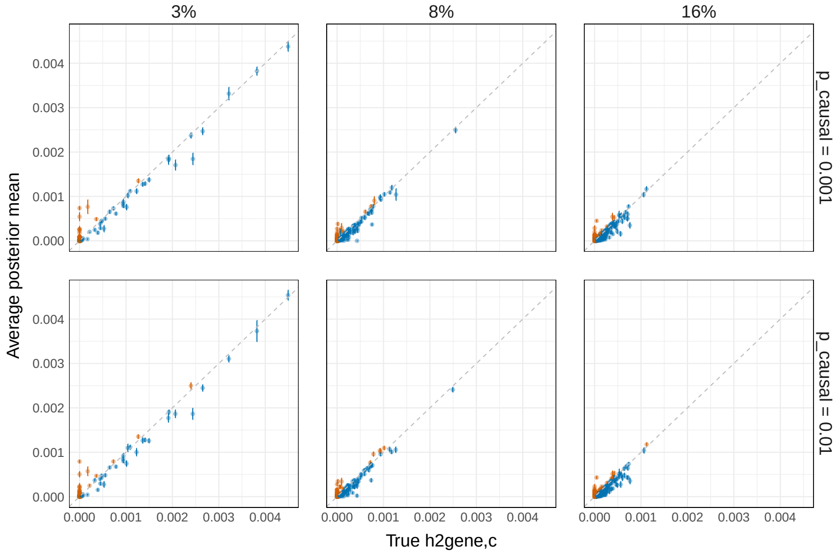


**Supplementary Figure 12**: Average estimated posterior mean of $h^2_{\mathrm{gene,c}}$ vs. true $h^2_{\mathrm{gene,c}}$ in simulations on chromosome 1 where 3%, 8%, or 16% of genes are causal. Orange points are genes with significant upward bias; blue points are all other genes. Bias is estimated as the average error from 30 simulation replicates and is considered "significant" if the error bars ($\pm 1.96$ s.e.m.) lie above the true value of $h^2_{\mathrm{gene,c}}$. Total $h^2_G = 0.05$, cumulative $h^2_{\mathrm{gene,t}} = 0.03$, MAF $>$ 0.005, N=291K.

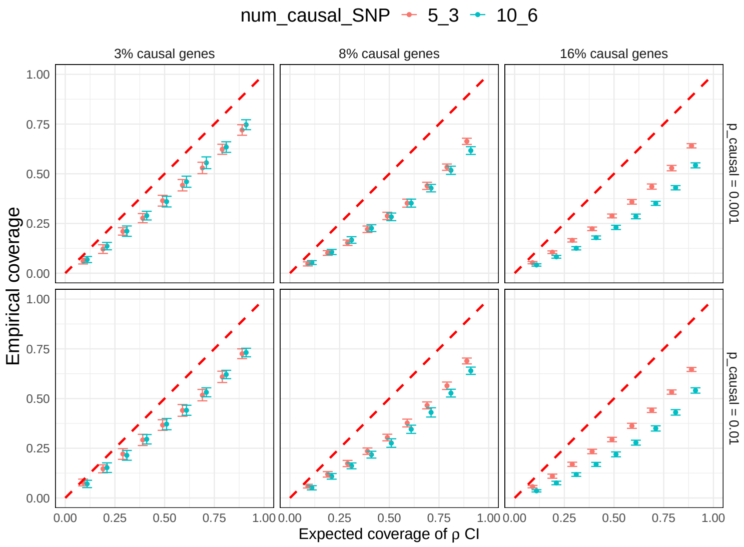


**Supplementary Figure 13**: Calibration of total $h^2_{\text{gene,t}}$ $\rho$-CIs for $\rho \in \{0.1, 0.2, ..., 0.9\}$. Empirical coverage for a given gene is the proportion of simulation replicates (out of 30) in which $\rho$-CI overlaps the true gene-level heritability. Causal genes contain either 5 (red) or 10 (blue) causal variants; their respective TSSs contain either 3 (red) or 6 (blue) causal variants (Methods). Chromosome 1, MAF $>$ 0.005, N=291K, 1,083 genes.

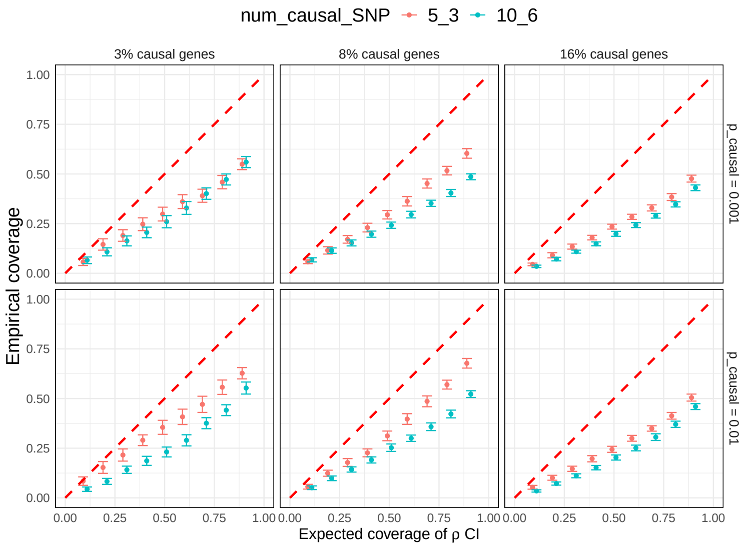


**Supplementary Figure 14**: Calibration of $h^2_{\text{gene,r}}$ $\rho$-CIs for $\rho \in \{0.1, 0.2, ..., 0.9\}$. Empirical coverage for a given gene is the proportion of simulation replicates (out of 30) in which $\rho$-CI overlaps the true gene-level heritability. Causal genes contain either 5 (red) or 10 (blue) causal variants; their respective TSSs contain either 3 (red) or 6 (blue) causal variants (Methods). Chromosome 1, MAF $>$ 0.005, N=291K, 1,083 genes.

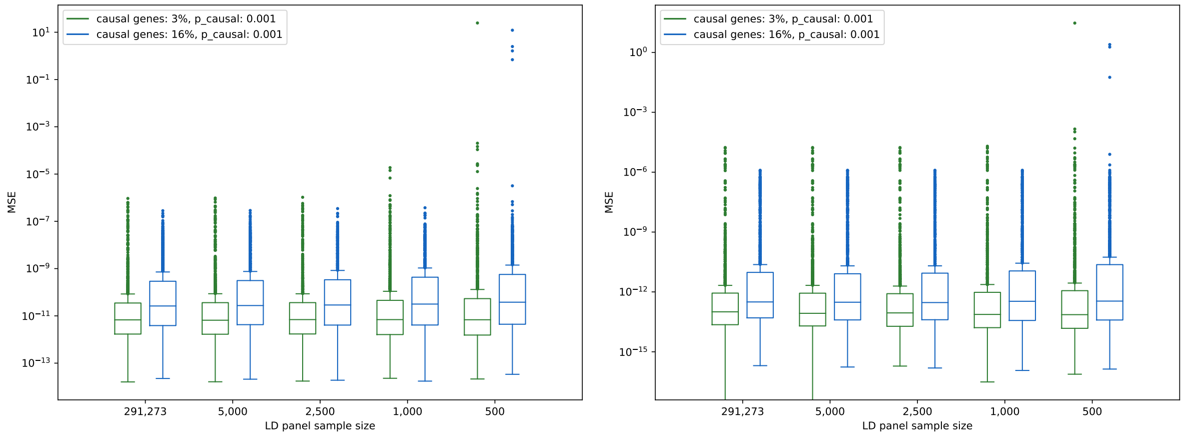


**Supplementary Figure 15**: MSE of $\hat{h}^2_{\text{gene,t}}$ (left) and $\hat{h}^2_{\text{gene,r}}$ (right) with respect to LD panel sample size (x-axis) in simulations (chromosome 1, MAF $>$ 0.005, N=290K, 1,083 genes). (Note: y-axes are different).

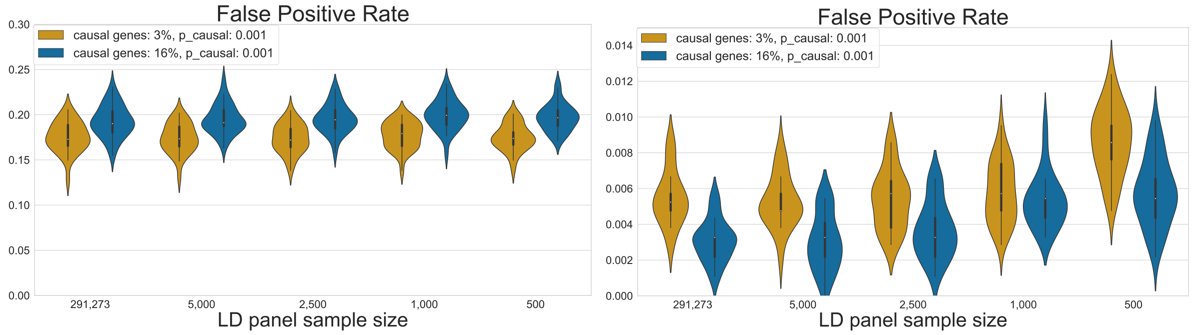


**Supplementary Figure 16**: False positive rate (FPR) with respect to LD panel sample size in simulations. Here, FPR is defined as the proportion of genes with 90%-CIs $>$ 0 that are noncausal. Left: $h^2_{\text{gene,t}} = 0$. Right: $h^2_{\text{gene,r}} = 0$. $\rho = 0.9$, chromosome 1, MAF $>$ 0.005, N=290K, 1,083 genes. LD panels were generated by sampling individuals from the GWAS cohort. (Note: y-axes are different.)

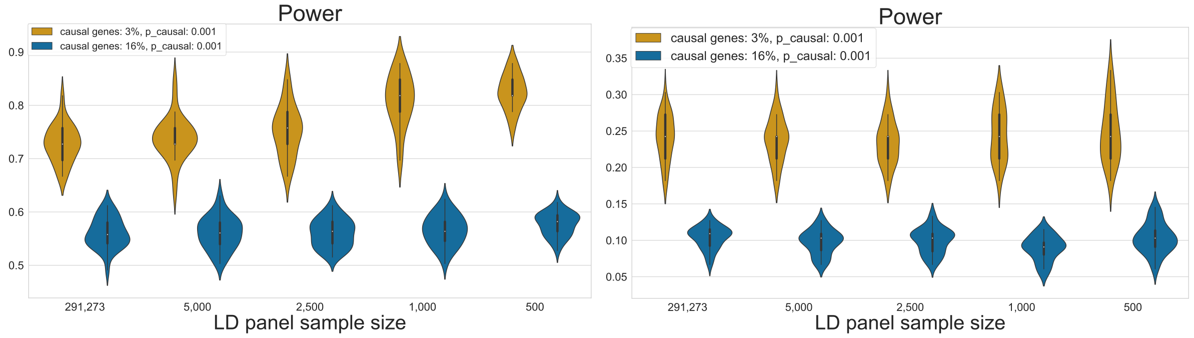


**Supplementary Figure 17**: Power with respect to LD panel sample size in simulations. Here, power is defined as the proportion of true causal genes that are correctly identified as causal using the cutoff 90%-CIs $>$ 0. Left: $h^2_{\text{gene,t}} > 0$. Right: $h^2_{\text{gene,r}} > 0$. $\rho = 0.9$, chromosome 1, MAF $>$ 0.005, N=290K, 1,083 genes. LD panels were generated by sampling individuals from the GWAS cohort. (Note: y-axes are different.)

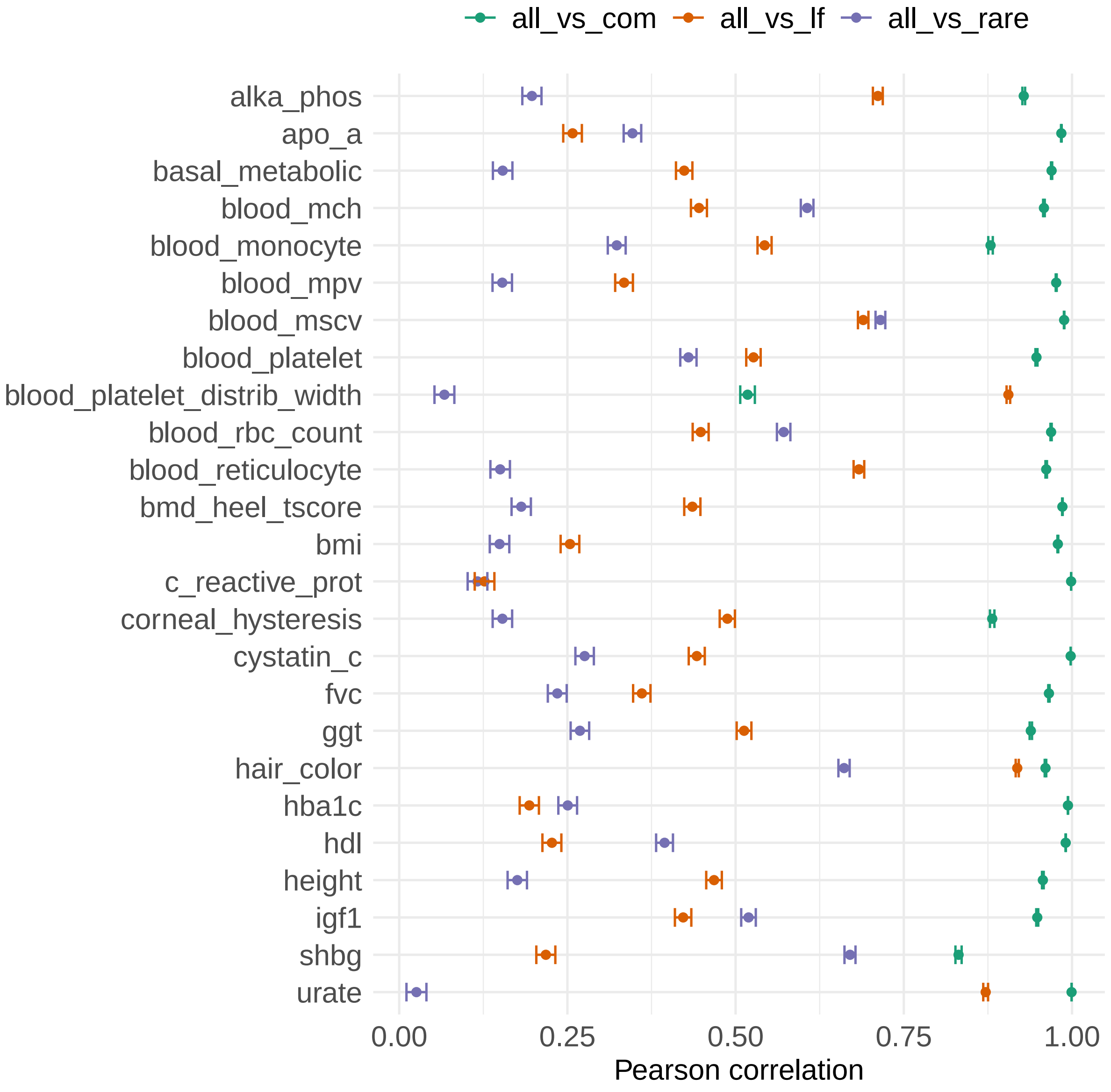


**Supplementary Figure 18**: Correlation of $\hat{h}^2_{\text{gene,t}}$ with $\hat{h}^2_{\text{gene,c}}$ (green), $\hat{h}^2_{\text{gene,lf}}$ (orange), and $\hat{h}^2_{\text{gene,r}}$ (purple) for 25 UK Biobank traits. Error bars mark 95% confidence intervals.

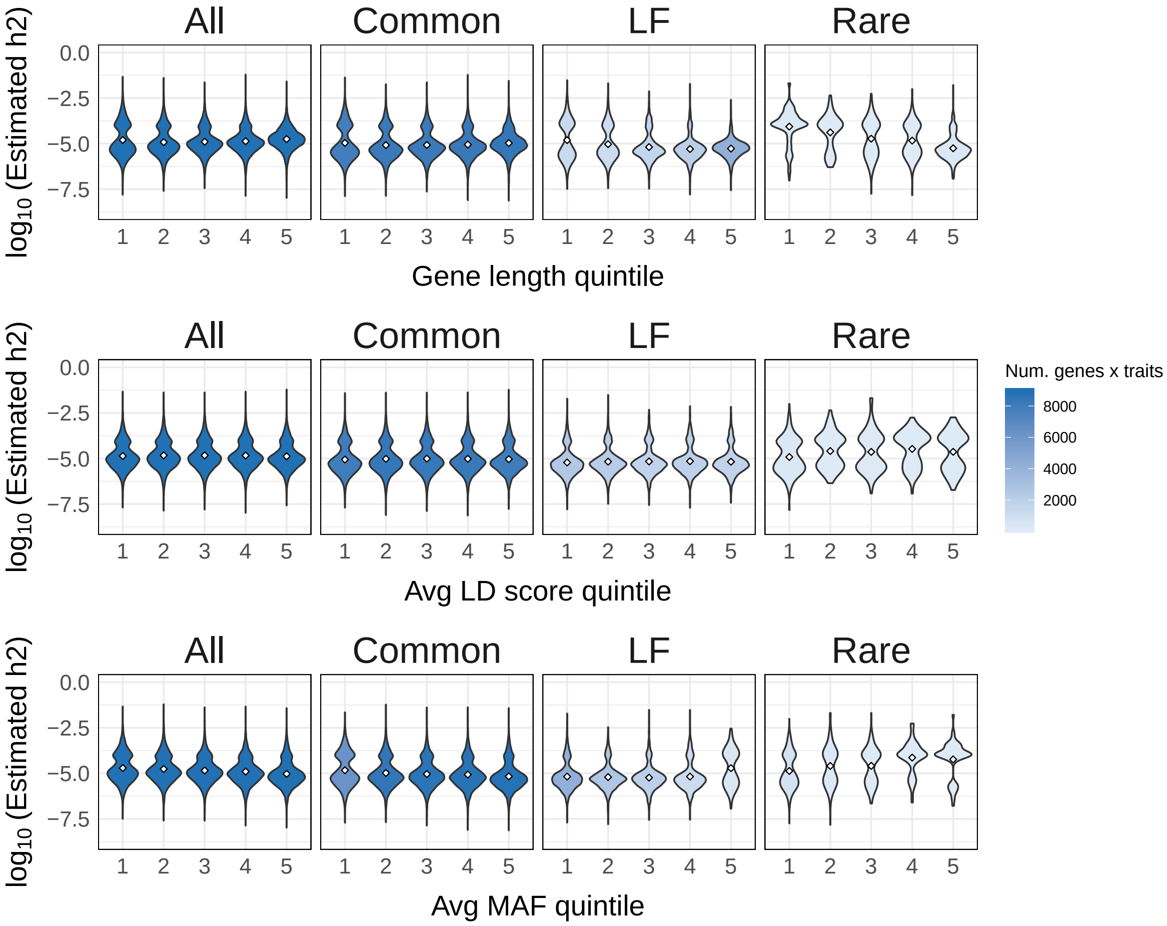


**Supplementary Figure 19**: Distributions of $\hat{h}^2$ for nonzero-heritability genes with respect to gene length (top), average LD score of variants assigned to gene (middle), average MAF of variants assigned to gene (bottom). Each point in each violin plot is an estimate for a unique gene-trait pair (25 traits in total). Violin plots are shaded to indicate the number of data points in the distribution. Diamonds mark the means of the distributions.

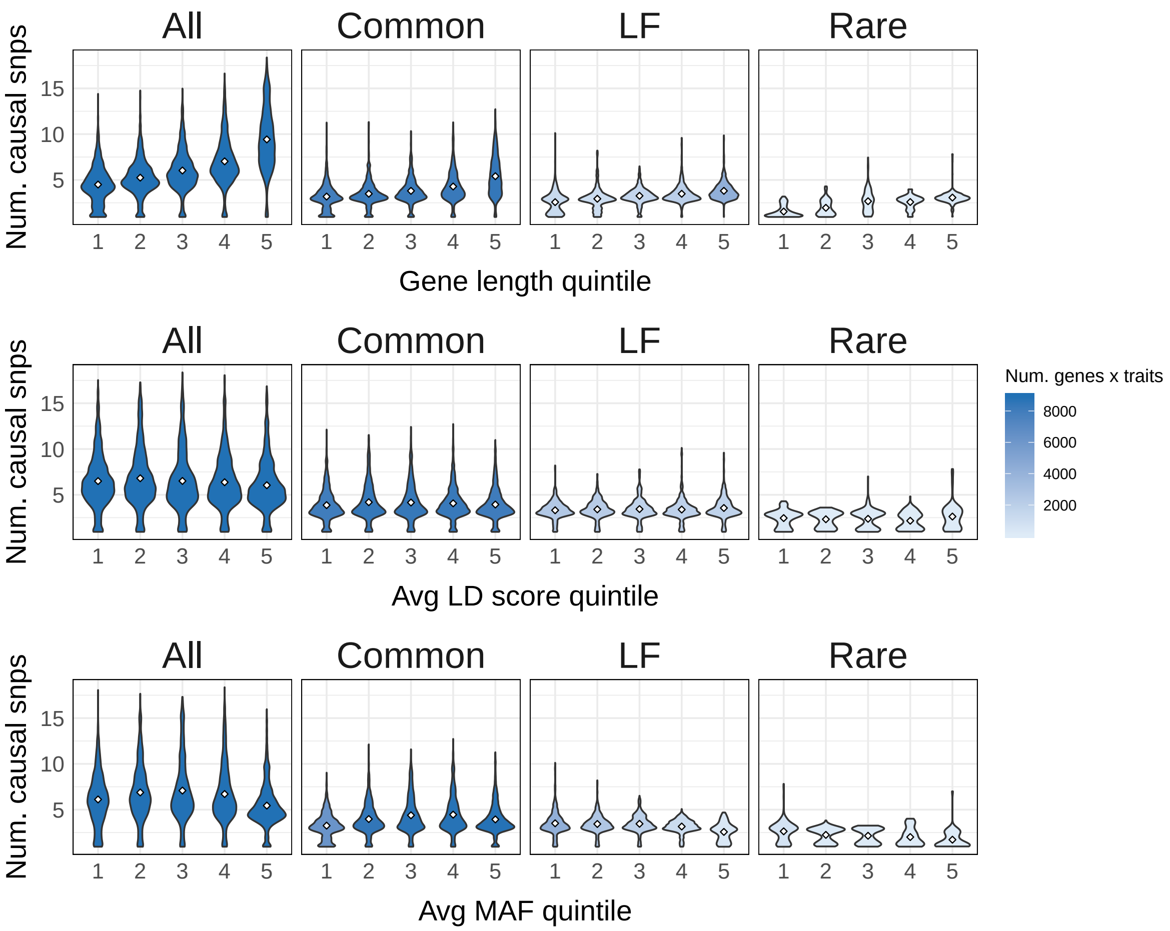


**Supplementary Figure 20**: Distributions of estimated number of causal variants in nonzero-heritability genes with respect to gene length (top), average LD score of variants assigned to gene (middle), and average MAF of variants assigned to gene (bottom). Violin plots are shaded to indicate the number of data points in the distribution. Each point in each violin plot is an estimate for a unique gene-trait pair (25 traits in total). Diamonds mark the means of the distributions.

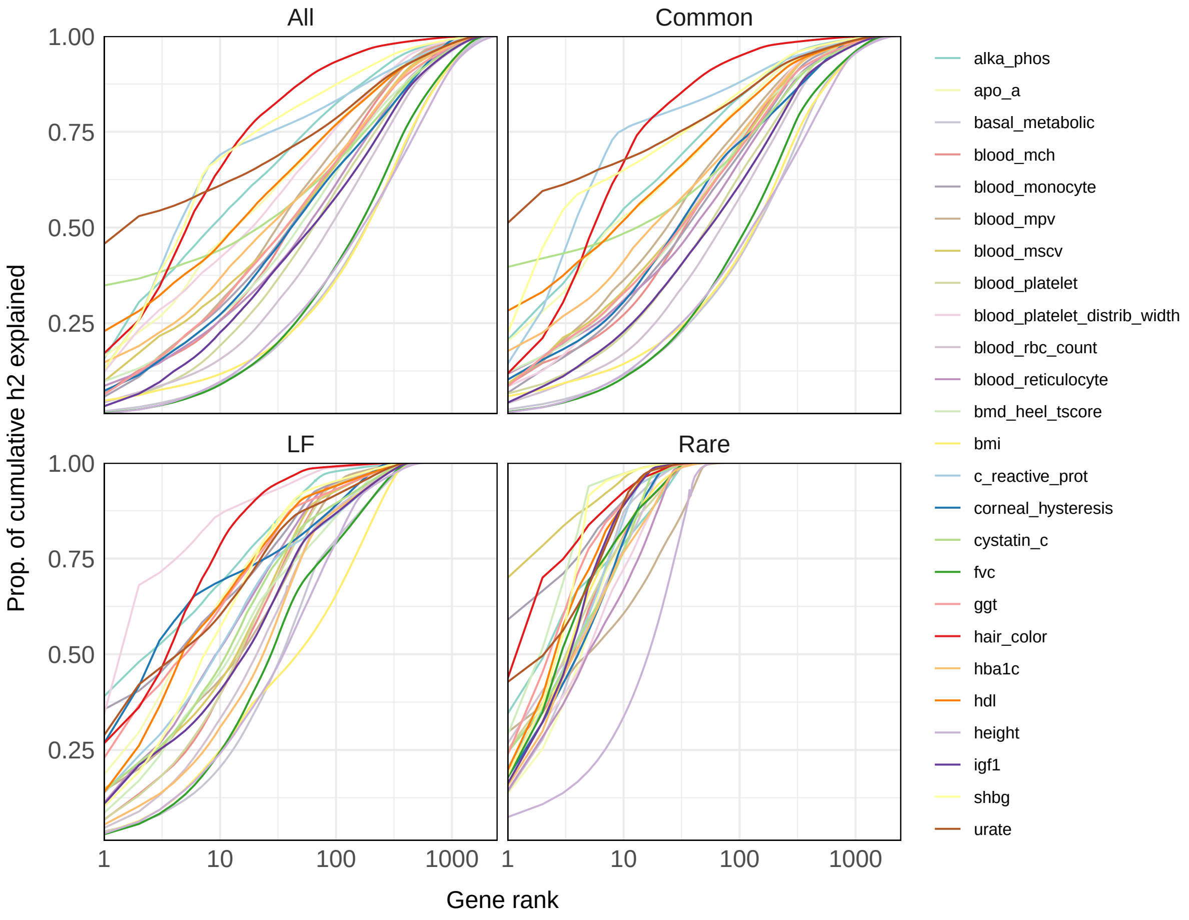


**Supplementary Figure 21**: Empirical cumulative distribution cumulative heritability for 25 traits. Each curve can be read as, "the top X genes explain Y% of the cumulative gene-level heritability for a given trait." Cumulative heritability is estimated as the summation of posterior mean estimates for nonzero-heritability genes (90%-CI $>$ 0). Clockwise from the top left: $\hat{h}^2_{\text{gene,t}}$, $\hat{h}^2_{\text{gene,c}}$, $\hat{h}^2_{\text{gene,r}}$, and $\hat{h}^2_{\text{gene,lf}}$.

| % causal genes | $p_{\mathrm{causal}}$ | $\rho$ | Underestimated | | Overestimated | |
|---|---|---|---|---|---|---|
| | | | Avg num genes | Avg % | Avg num genes | Avg % |
| 3% | 0.001 | 0.90 | 6.55 (0.28) | 19.29 (0.82) | 2.50 (0.13) | 7.37 (0.40) |
| 3% | 0.001 | 0.95 | 4.82 (0.26) | 14.17 (0.78) | 1.88 (0.13) | 5.58 (0.40) |
| 3% | 0.01 | 0.90 | 6.25 (0.26) | 18.42 (0.79) | 2.98 (0.17) | 8.74 (0.48) |
| 3% | 0.01 | 0.95 | 4.68 (0.23) | 13.80 (0.68) | 2.18 (0.14) | 6.43 (0.41) |
| 8% | 0.001 | 0.90 | 25.47 (0.64) | 29.39 (0.70) | 5.72 (0.24) | 6.62 (0.28) |
| 8% | 0.001 | 0.95 | 19.77 (0.63) | 22.80 (0.70) | 4.40 (0.20) | 5.09 (0.23) |
| 8% | 0.01 | 0.90 | 23.13 (0.64) | 26.70 (0.70) | 5.93 (0.23) | 6.86 (0.27) |
| 8% | 0.01 | 0.95 | 17.60 (0.58) | 20.30 (0.65) | 4.48 (0.24) | 5.18 (0.27) |
| 16% | 0.001 | 0.90 | 61.82 (1.43) | 36.22 (0.71) | 7.87 (0.28) | 4.62 (0.16) |
| 16% | 0.001 | 0.95 | 49.62 (1.38) | 29.05 (0.71) | 5.55 (0.27) | 3.26 (0.15) |
| 16% | 0.01 | 0.90 | 60.55 (1.58) | 35.46 (0.81) | 8.83 (0.31) | 5.20 (0.18) |
| 16% | 0.01 | 0.95 | 48.95 (1.47) | 28.64 (0.76) | 6.20 (0.27) | 3.66 (0.16) |

**Supplementary Table 1.** Calibration of h2gene $\rho$-CIs with respect to the number of causal genes, proportion of causal variants ($p_{\mathrm{causal}}$), and $\rho \in \{0.90, 0.95\}$ in simulations (chromosome 1, MAF > 0.5%, 1,083 protein-coding genes, cumulative $h^2_{\mathrm{gene}} = 0.03$ ). "Underestimated" and "overestimated" refer to genes whose $\rho$-CIs lie below and above their true gene-level heritability, respectively. For each simulation setup, we report the average (and s.e.m.) of the number and percentage of underestimated/overestimated genes in 30 simulation replicates.

| % causal genes | $p_{causal}$ | $\rho$ | Underestimated | | Overestimated | |
|---|---|---|---|---|---|---|
| | | | Avg num genes | Avg % | Avg num genes | Avg % |
| 3% | 0.001 | 0.90 | 9.70 (0.38) | 42.41 (0.93) | 0.52 (0.09) | 2.15 (0.38) |
| 3% | 0.001 | 0.95 | 8.67 (0.35) | 37.88 (0.95) | 0.35 (0.08) | 1.45 (0.31) |
| 3% | 0.01 | 0.90 | 8.98 (0.44) | 38.40 (0.97) | 0.67 (0.11) | 2.58 (0.40) |
| 3% | 0.01 | 0.95 | 7.80 (0.40) | 33.31 (1.00) | 0.43 (0.09) | 1.66 (0.34) |
| 8% | 0.001 | 0.90 | 25.82 (0.95) | 43.76 (0.98) | 1.07 (0.15) | 1.78 (0.25) |
| 8% | 0.001 | 0.95 | 22.73 (0.87) | 38.51 (0.95) | 0.52 (0.10) | 0.85 (0.16) |
| 8% | 0.01 | 0.90 | 22.97 (1.09) | 38.49 (1.26) | 0.87 (0.11) | 1.53 (0.21) |
| 8% | 0.01 | 0.95 | 19.52 (0.95) | 32.65 (1.12) | 0.55 (0.09) | 0.97 (0.16) |
| 16% | 0.001 | 0.90 | 61.03 (1.63) | 53.38 (0.65) | 1.35 (0.17) | 1.27 (0.16) |
| 16% | 0.001 | 0.95 | 53.90 (1.41) | 47.20 (0.60) | 0.68 (0.12) | 0.65 (0.12) |
| 16% | 0.01 | 0.90 | 57.87 (1.57) | 50.60 (0.65) | 1.27 (0.16) | 1.19 (0.16) |
| 16% | 0.01 | 0.95 | 51.08 (1.36) | 44.73 (0.59) | 0.78 (0.12) | 0.75 (0.12) |

**Supplementary Table 2.** Calibration of h2rare $\rho$-CIs with respect to the number of causal genes, proportion of causal variants ($p_{\text{causal}}$), and $\rho \in \{0.90, 0.95\}$ in simulations (chromosome 1, MAF > 0.5%, 1,083 protein-coding genes, cumulative $h^2_{\text{gene}} = 0.03$ ). "Underestimated" and "overestimated" refer to genes whose $\rho$-CIs lie below and above their true gene-level heritability, respectively. For each simulation setup, we report the average (and s.e.m.) of the number and percentage of underestimated/overestimated genes in 30 simulation replicates.
